# Paleogenomic Evidence for Genetic Heterogeneity and Prior Admixture in Gothic-Associated Communities of Late Antique Bulgaria

**DOI:** 10.64898/2026.03.03.709317

**Authors:** Svetoslav Stamov, Todor Chobanov, Tianyi Wang, Kremena Stoeva, Dimcho Momchilov, Andrey Aladzhov, Kaloyan Chobanov, Miroslav Klasnakov, Georgi Stamov, Milen Nikolov, Desislava Nesheva, Peter Heather, Draga I. Toncheva, Milen Zamfirov, Iosif Lazaridis, David Reich

## Abstract

We report genome-wide ancient DNA from 38 individuals associated with Gothic-period mortuary contexts at two Bulgarian sites: the Aquae Calidae necropolis (∼320 - 375 CE; n=23) and the Aul of Khan Omurtag (AKO; ∼350 - 489 CE; n=15). Although linked by closely similar Gothic-associated material culture and east - west Arian burial practices, the two assemblages are genetically distinct. Aquae Calidae individuals are modeled with predominantly Anatolian-related ancestry (50 - 85% in qpAdm), whereas AKO individuals are modeled with predominantly Wielbark/Chernyakhov-related northern ancestry (60 - 78%). Across the full assemblage, no single ancestry model fits all individuals, whose profiles range from nearly unadmixed Chernyakhov-like to predominantly Anatolian-related. DATES analysis places admixture between northern European and southern Balkan/Anatolian-related ancestry components approximately 11 - 13 generations before burial (point estimate ∼50 CE; 95% CI, 85 BCE - 183 CE; Z=5.26), raising the possibility that this mixture predated the earliest documented Gothic - Roman contacts (∼170 CE). The convergence of dates across alternative southern proxies, together with reduced fit when Balkan and Anatolian sources are pooled into a single southern reference, favors admixture involving a pre-blended Balkan - Anatolian substrate over two temporally distinct southern admixture pulses. This signal was not recovered in the non-Gothic Roman-period Balkan populations tested as controls and is consistent with admixture in a trans-Danubian frontier setting, potentially including Roman Dacia after 106 CE. These findings support models in which Gothic affiliation in the Balkans operated as a cultural-political framework encompassing populations of diverse biological ancestry.

## 1. Introduction

The Goths represent one of the most consequential groups of the Migration Period (c. 300 - 700 CE). Historical sources trace their origins to the Wielbark culture of northern Poland^1^ (1st - 4th centuries CE), characterized by Scandinavian-related ancestry and elevated Y-haplogroup I1 frequencies (∼41%)^2^, followed by incorporation into the multi-ethnic Chernyakhov culture (late 2nd - 5th centuries CE), where Gothic-associated populations merged with Dacians, Sarmatians, and Alans^3^. The 382 CE foederati treaty under Theodosius I established Gothic communities in the Balkans; Theodoric’s 488 CE departure for Italy left residual populations in the region^4^. Detailed chronology and site archaeology are in Supplementary Note S4.

Essentialist models treat "Goths" as a biologically coherent migrating group ^5^; ethnogenesis models treat Gothic identity as a cultural-political construct maintained by shared institutions rather than biological descent ^6,7^. Ancient DNA can constrain competing models of ethnogenesis by quantifying ancestry patterns, admixture timing, and kinship structure relative to cultural attribution.

### Study Sites

Aquae Calidae (Thrace, ∼320 - 375 CE, n=23) is a 4th-century necropolis adjacent to Roman thermal baths. Burials ignore prior Roman structures; some individuals exhibit artificial cranial deformation^8^. Extreme male bias (17:3 among sexed individuals) is consistent with a military context. Diagnostic Gothic artifacts (fibulae, beads) and Arian burial orientations are present. Jordanes (Getica 20.109)^9^ places Gothic arrival ∼270 CE; Procopius (On Buildings 3.7.18 - 23)^10^ records "barbarian villagers" occupying the site until Justinian’s expulsion.

The Aul of Khan Omurtag (AKO, Moesia Secunda, ∼350 - 489 CE, n=15) is an Arian episcopal center with four basilicas, potentially tied to Ulfilas’ 348 CE settlement of Christian Goths in Moesia^11^. Sequential necropoleis (C1 - C5) span pre-Hunnic (C1/C2), a transitional phase (C3) whose burials may be contemporaneous with the C4 basilica despite their earlier stratigraphy, Hunnic-period contexts (C4), and post-Nedao occupation (C5). No C3 individuals yielded usable genomic data. Phase definitions and dating anchors are in Supplementary Note S4.

**Figure 1.**
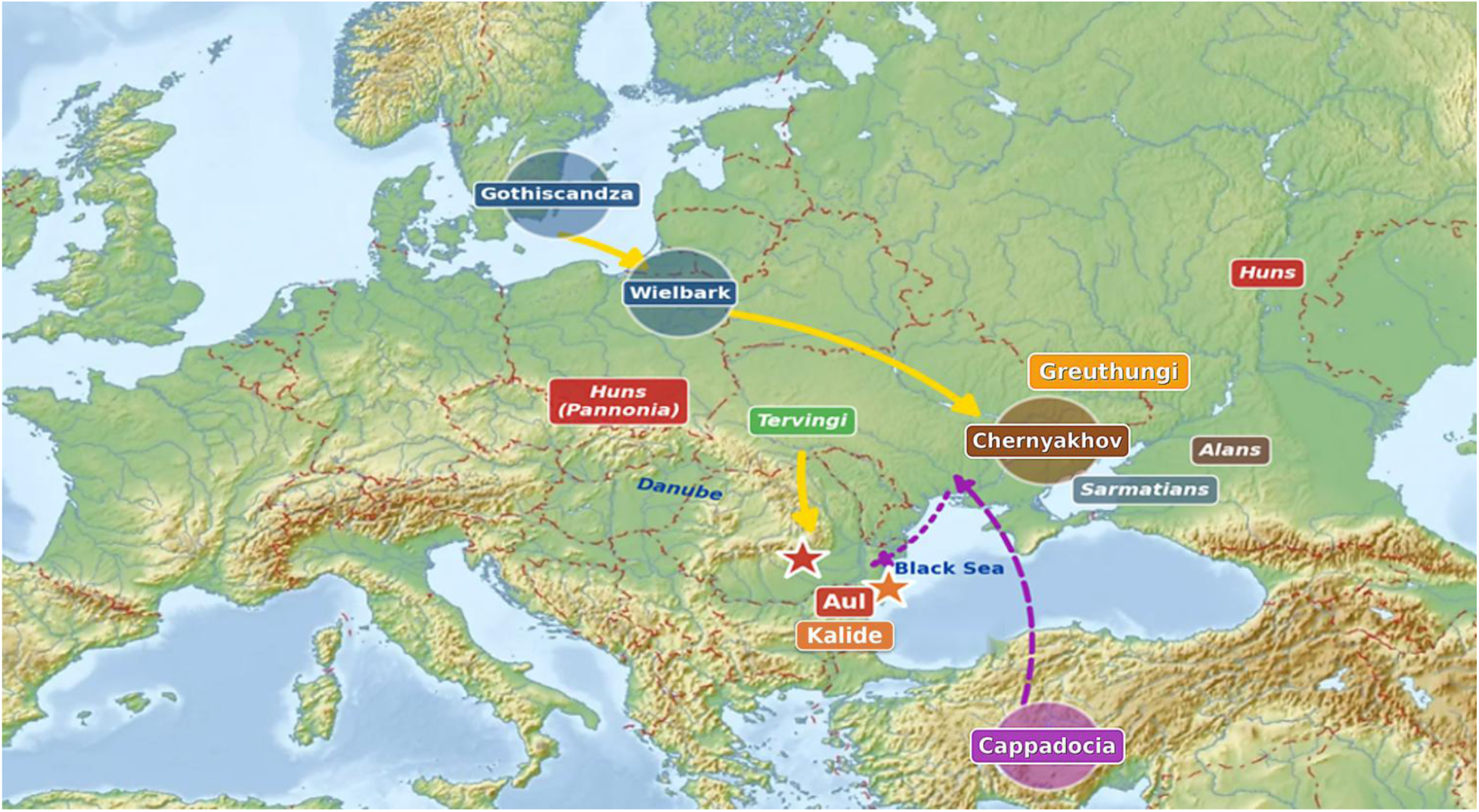
Site locations and inferred Gothic migration routes (320 - 490 CE). Aquae Calidae (Thrace) and the Aul of Khan Omurtag (Moesia Secunda) are shown with major Gothic movement corridors referenced in the text (after ^12^,^13^).

## 2. Results

### 2.1 The Two Sites Form Genetically Distinct Clusters

PCA projected onto modern West Eurasian variation (PC1 - 2 capturing ∼19.9% and ∼9.9% of variance) reveals pronounced differentiation between the two sites. The two assemblages sharing Gothic material culture and Arian burial rites are genetically more distant from each other than either is from its respective closest reference population: AKO individuals (red triangles) cluster with Iron Age Wielbark and Chernyakhov Migration Period groups in the northern portion of the plot, while Aquae Calidae individuals separate into two subclusters - a southern group (blue diamonds, n=12) shifted toward Ancient Anatolian and Byzantine Marmara references, and a northern group (blue squares, n=10) aligned with Byzantine Marmara/Iznik populations - with Bulgaria Late Antiquity (green circles) ranking as the top f3 affinity for Aquae Calidae as a whole (rank #1) but a much weaker signal at AKO (rank #21). One individual (I40570, asterisk) positions near ancient Levantine and Egyptian references, separated from all other Aquae Calidae individuals by a gap exceeding the full within-site PC2 range. Together, these patterns are inconsistent with a single biological profile underlying Gothic cultural affiliation at either site.

**Figure 2.**
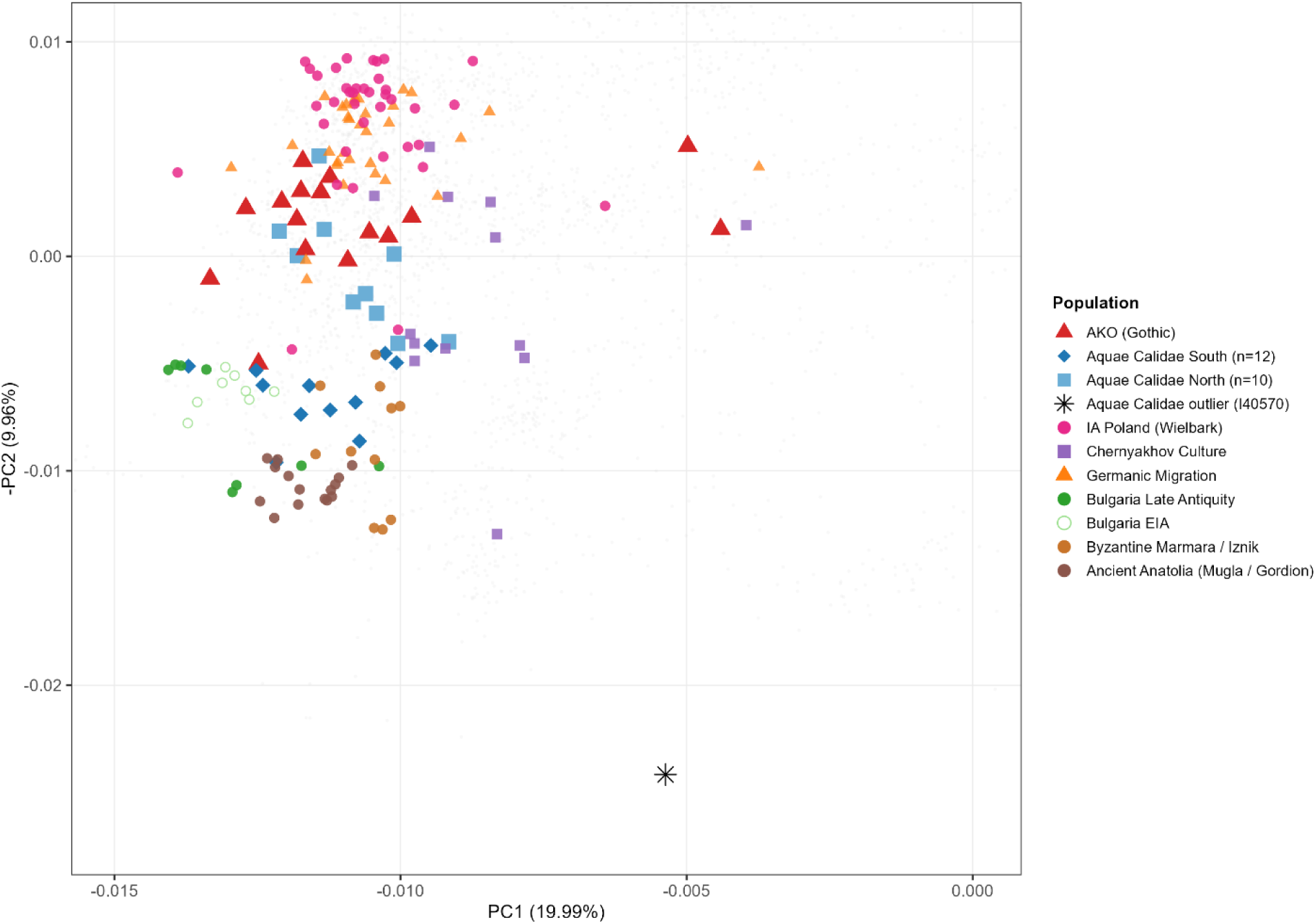
PCA of Gothic-associated individuals from Aquae Calidae and AKO projected onto modern West Eurasian variation (PC1 - 2 capturing 19.99% and 9.96% of variance).

Outgroup f3 statistics quantify the divergence: Aquae Calidae ranks Bulgaria_LAntiquity as its top affinity (f3=0.0524, rank #1), a signal 2 - 3× stronger than at AKO (rank #21), reflecting greater local Balkan continuity or pre-arrival Mediterranean admixture. Affinities with eastern nomadic groups carrying substantial East Asian ancestry are uniformly low at both sites (f3 < 0.040), consistent with limited East Asian-related ancestry in the main assemblage. Full f3 results are in Supplementary Table S1 and in Supplementary_Note_S5_f3.

**Figure 3.**
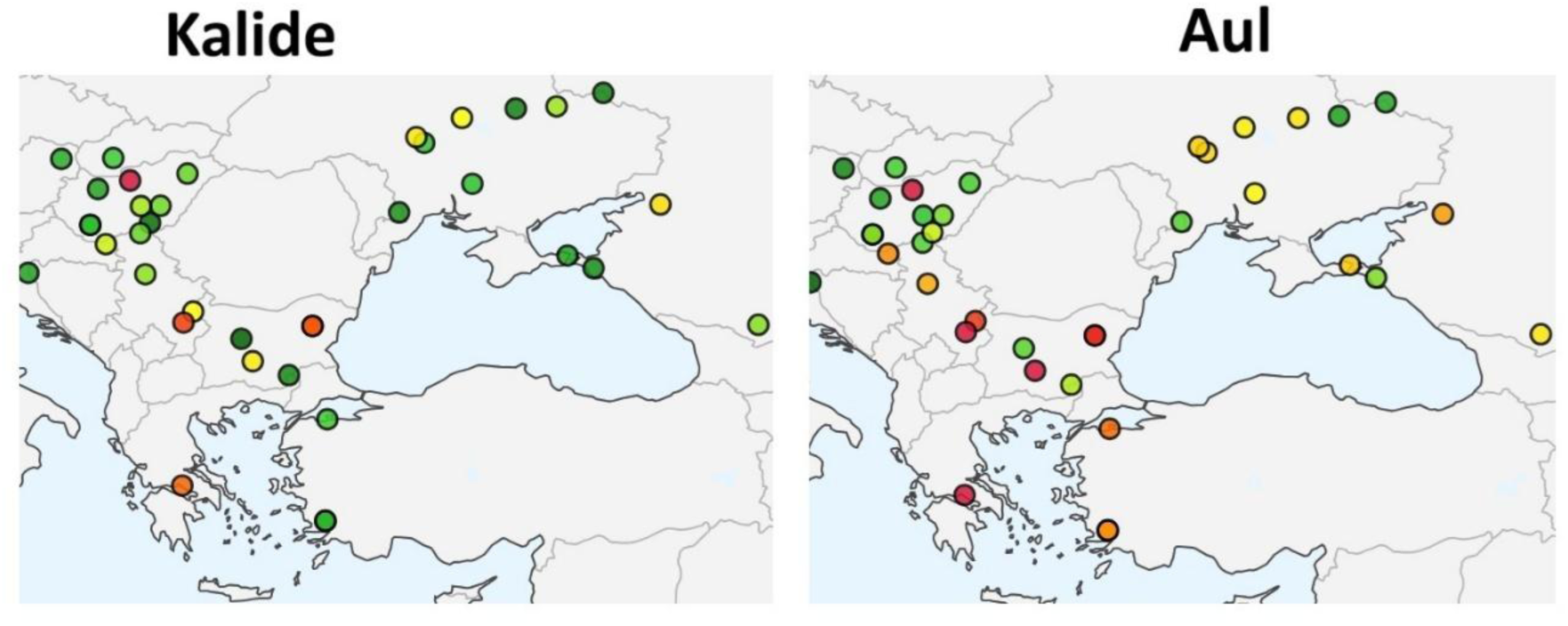
Outgroup f3 statistics heatmaps for Aquae Calidae (top) and AKO (bottom). Form: f3(Mbuti; Target, Reference). Color gradient from green (higher affinity) to red (lower affinity). Aquae Calidae ranks Bulgaria_LAntiquity as its top affinity (rank #1); AKO ranks it #21, reflecting distinct ancestry profiles across the two sites despite shared material culture.

### 2.2 Ancestry Is Predominantly Anatolian at Aquae Calidae and Pontic at AKO

We applied qpAdm admixture modeling using both proximal (temporally relevant) and distal source sets (see Methods; full outputs in Supplementary Tables S2 - S4).

#### Aquae Calidae

The southern subcluster (PCA-South, n=12) is consistent with ∼48% Central Anatolian (Kalehöyük ∼2000 BCE) + ∼34% Bulgaria_EIA + ∼18% Chernyakhov (p=0.713; Table 1). Models excluding Anatolian-related proxies are strongly rejected (p < 10⁻¹⁴⁹); Wielbark-only models fail (p < 10⁻²¹⁶). We tested ∼100 ancient Anatolian samples from >20 populations spanning Roman, Byzantine, Iron Age, and Bronze Age periods; Kalehöyük provides the best-fitting currently available proxy for the Anatolian-related component in Aquae Calidae, in the absence of published Roman-era central Anatolian reference data. The northern subcluster (PCA-North, n=10) requires a distinct source: no feasible models accept Kalehöyük, while Roman-Byzantine Marmara/Iznik fits well (best model: ∼43% Iznik_Byzantine + ∼27% Chernyakhov + ∼30% Chernyakhov_o; p=0.805; Table 2), suggesting temporally or geographically distinct incorporation pathways within the same site.

**Table 1.** qpAdm results for Aquae Calidae South (n=12). Best - fitting feasible models. SE = standard error from jackknife resampling.

| Rank | Sources | Proportions (±SE) | p - value |
| --- | --- | --- | --- |
| 1 | Kalehöyük + Bulgaria_EIA + Wielbark | 54.0±5.2% + 26.5±4.8% + 19.5±3.9% | 0.865 |
| 2 | Kalehöyük + Bulgaria_EIA | 36.6±6.1% + 26.6±6.1% | 0.863 |
| 3 | Kalehöyük + Chernyakhov + Bulgaria_EIA | 47.9±5.5% + 18.1±4.2% + 34.0±5.1% | 0.713 |
| FAIL | Wielbark only | 100% | 10<sup>-202</sup> |

**Table 2.** qpAdm results for Aquae Calidae North (n=10). Note: no feasible models with Kalehöyük source (distinct from South).

| Rank | Sources | Proportions (±SE) | p - value |
| --- | --- | --- | --- |
| 1 | Iznik_Byzantine + Chernyakhov + Chernyakhov_o | 43.4±5.8% + 26.9±4.5% + 29.8±5.3% | 0.805 |
| 2 | Iznik_Byzantine + Chernyakhov + Chernyakhov_o + Wielbark | 43.4±5.9% + 24.2±4.7% + 29.5±5.4% + 2.9±3.1% | 0.755 |
| FAIL | Chernyakhov_o only | 100% | 0.008 |

#### AKO

Population-level proximal qpAdm models AKO as 60 - 78% Chernyakhov (or equivalent Wielbark rotations) + 15 - 30% Anatolian/Near Eastern + 5 - 15% steppe (p=0.479 - 0.963). Distal modeling documents progressive transformation across phases: EHG (a diagnostic proxy for northern European heritage) declines from 27.9±3.2% in C1 (∼350 CE) to 4.6 - 16.2±4.1% in C5 (∼454 - 489 CE), while Anatolian Neolithic Farmer ancestry (Turkey_N) rises from ∼21% to ∼32 - 50%, possibly suggesting continuous local admixture over ∼150 years. Individual-level model outputs by phase are in Supplementary Table S3.

**Table 3.**
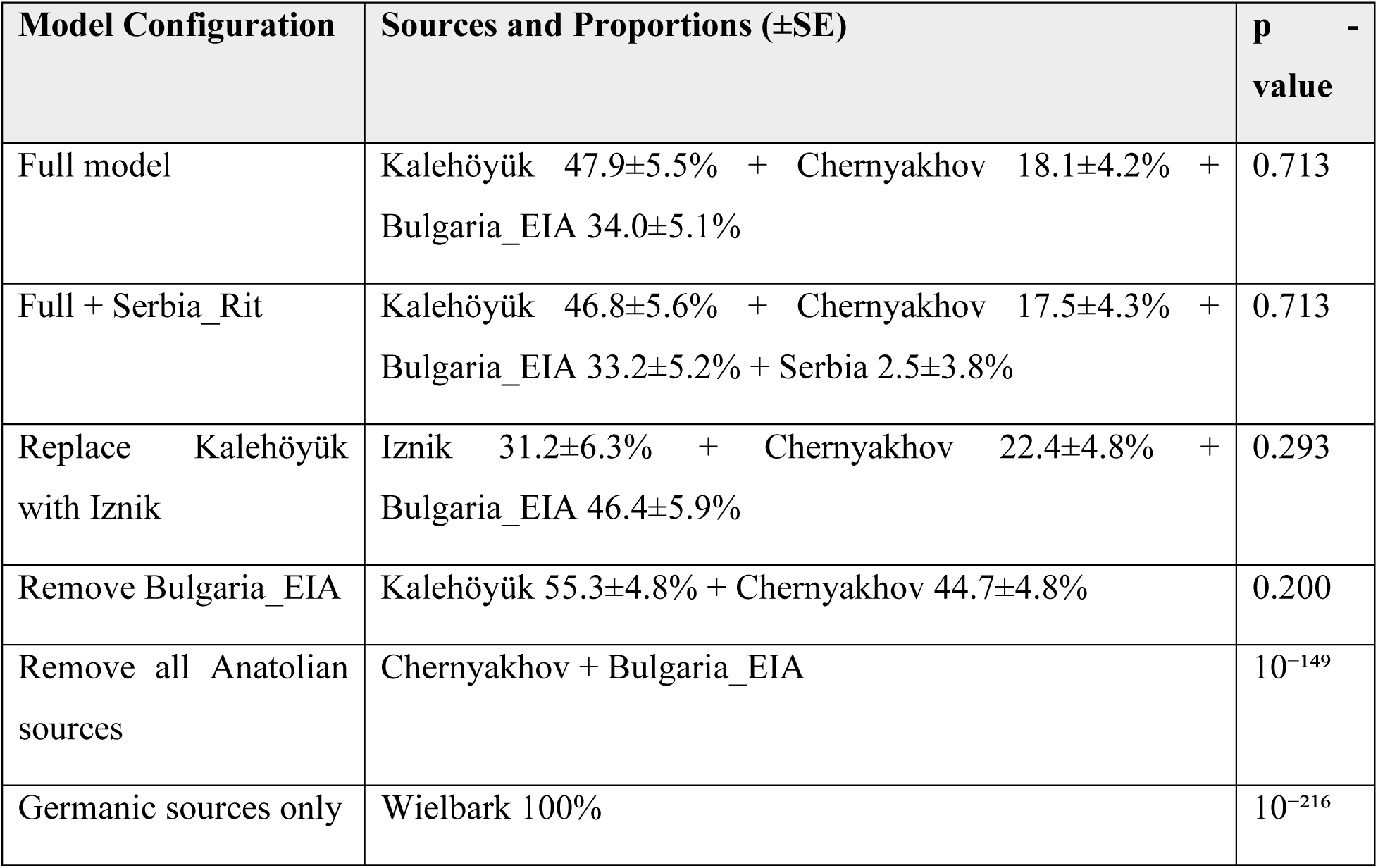
Model sensitivity analysis for Aquae Calidae South. Models (left + right), proportions, standard errors, and p - values.

| Model Configuration | Sources and Proportions (±SE) | p - value |
| --- | --- | --- |
| Full model | Kalehöyük 47.9±5.5% + Chernyakhov 18.1±4.2% + Bulgaria_EIA 34.0±5.1% | 0.713 |
| Full + Serbia_Rit | Kalehöyük 46.8±5.6% + Chernyakhov 17.5±4.3% + Bulgaria_EIA 33.2±5.2% + Serbia 2.5±3.8% | 0.713 |
| Replace Kalehöyük with Iznik | Iznik 31.2±6.3% + Chernyakhov 22.4±4.8% + Bulgaria_EIA 46.4±5.9% | 0.293 |
| Remove Bulgaria_EIA | Kalehöyük 55.3±4.8% + Chernyakhov 44.7±4.8% | 0.200 |
| Remove all Anatolian sources | Chernyakhov + Bulgaria_EIA | 10<sup>-149</sup> |
| Germanic sources only | Wielbark 100% | 10<sup>-216</sup> |

f4 statistics [f4(Mbuti, Anatolian; Aquae Calidae, Chernyakhov)] confirm excess Anatolian affinity in PCA-South individuals beyond the Chernyakhov baseline (Gordion Z=−3.11; Kalehöyük Z=−2.09; Mardin Z=−2.61), while PCA-North shows weaker signals (Z=−0.65 to −1.72). Full f4 results are in Supplementary Table S6.

**Figure 4A.**
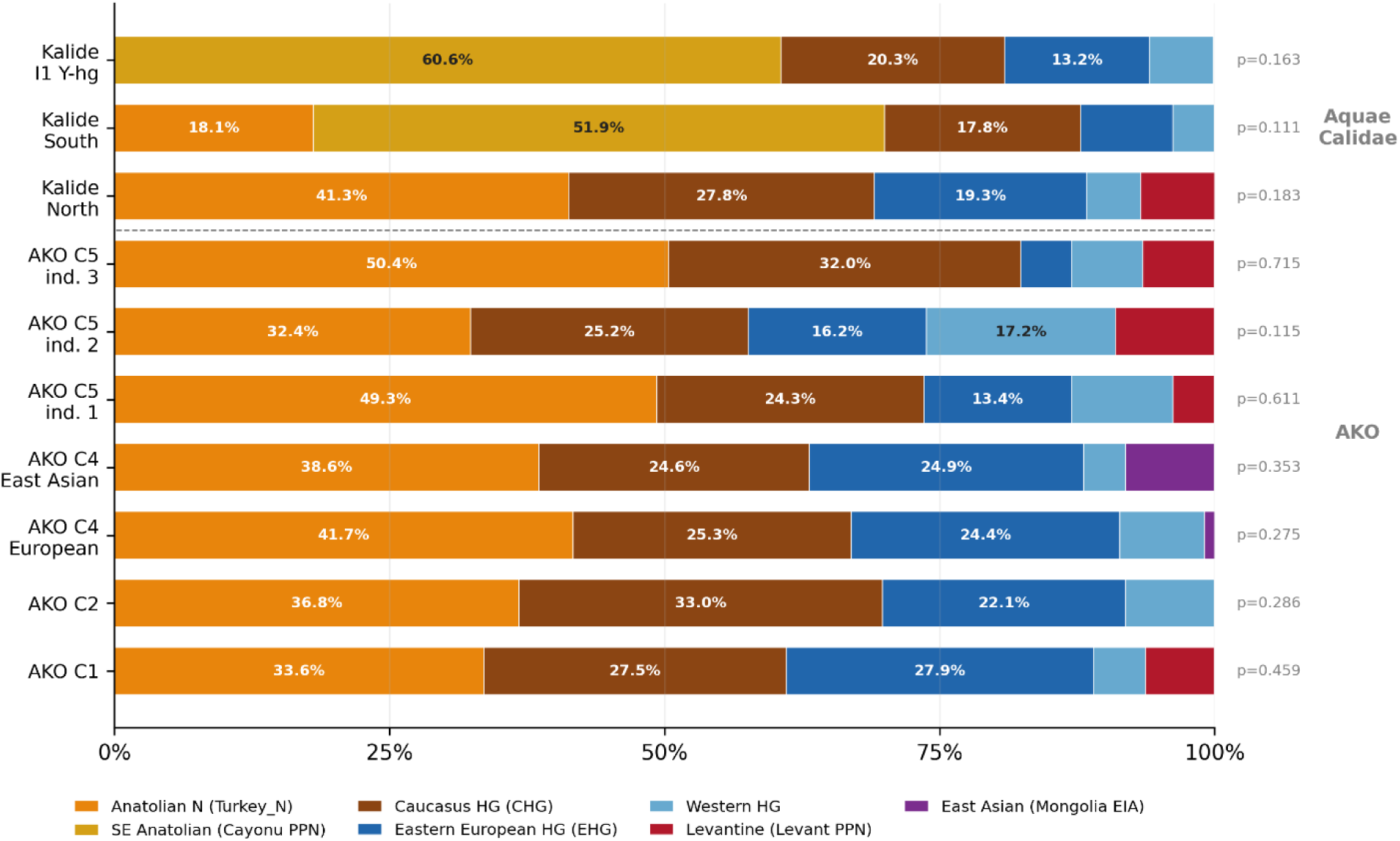
Distal qpAdm deep-ancestry decomposition using extended source set (Turkey_N, Cayonu_PPN, CHG, EHG, WHG, Levant_PPN, Mongolia_EIA). Aquae Calidae populations show dominant Anatolian Neolithic (Turkey_N) ancestry; AKO populations display more balanced profiles with substantial EHG components. All models pass at p > 0.05.

**Figure 4B.**
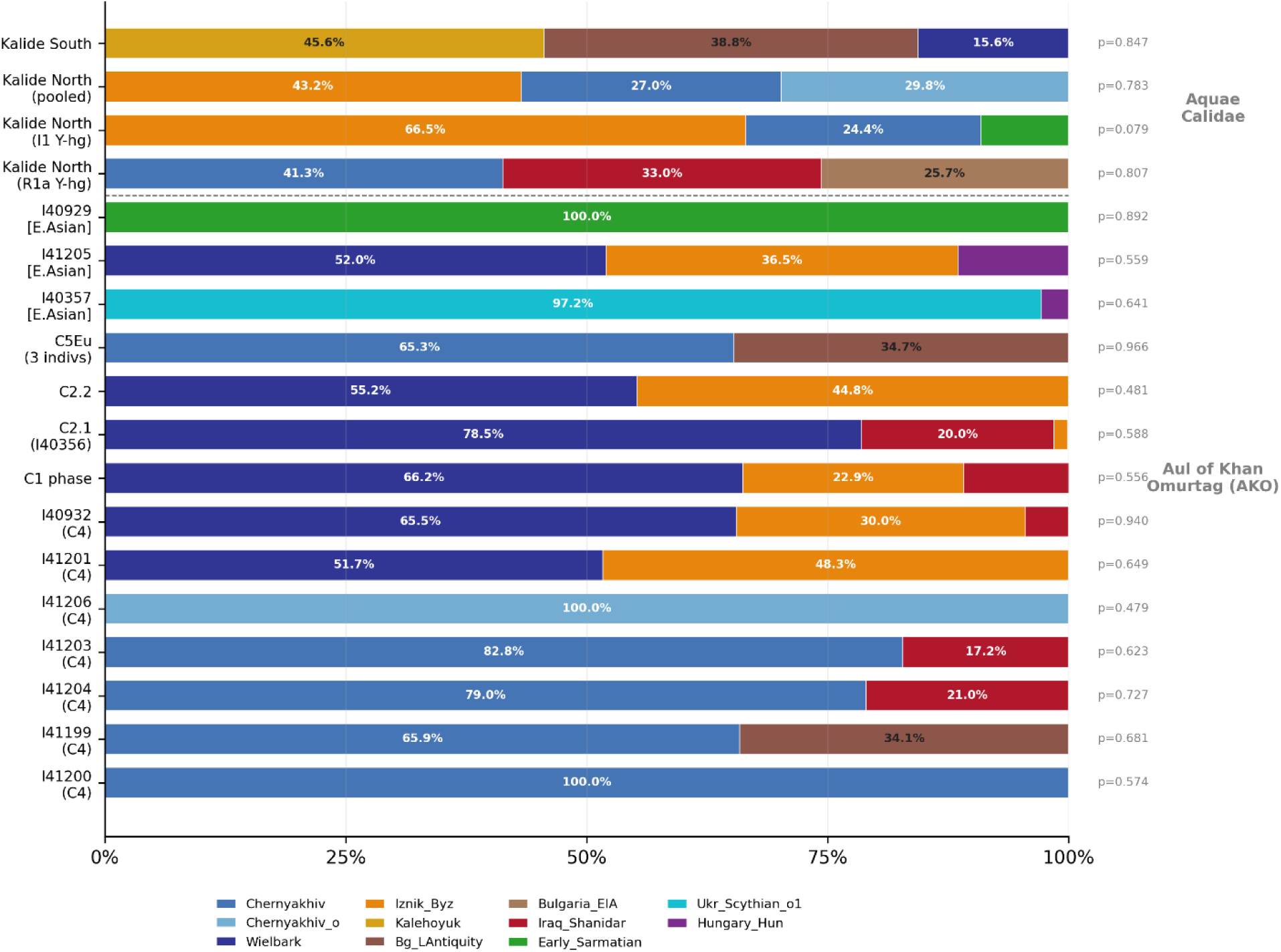
Proximal qpAdm ancestry models using temporally relevant source populations (Tables 1 - 3). Upper panel: Aquae Calidae subclusters (sources: Chernyakhov, Iznik_Byzantine, Kalehöyük, Bulgaria_EIA). Lower panel: AKO individuals and phase groups (sources: Chernyakhov, Wielbark, steppe-related proxies). All displayed models have p ≥ 0.05.

**Figure 4C.**
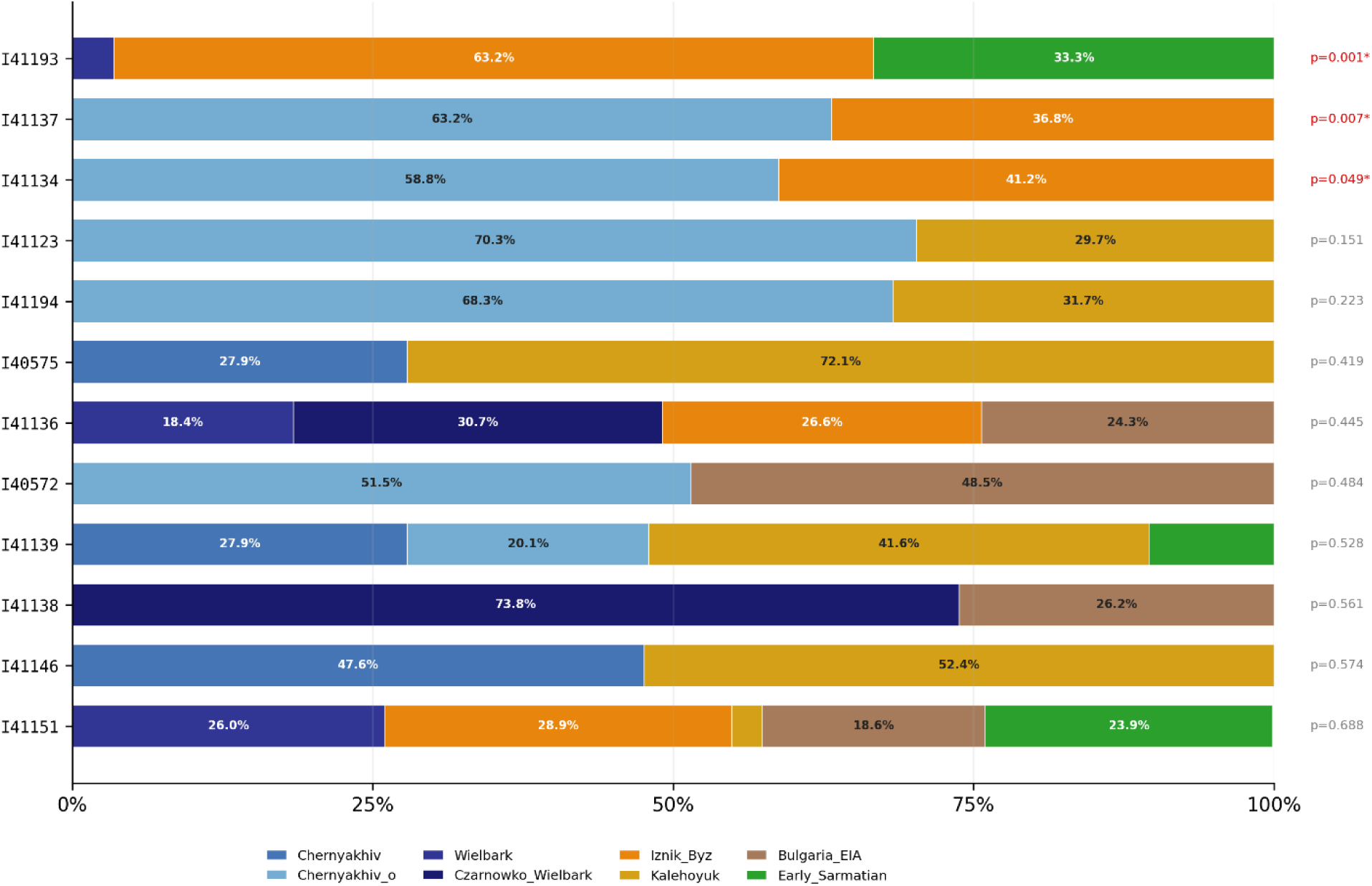
Individual-level proximal qpAdm models for the Aquae Calidae North subcluster (n=13). Each bar shows the best-fitting model per individual sorted by p-value; models rejected at p < 0.05 are marked with asterisks.

**Figure 4D.**
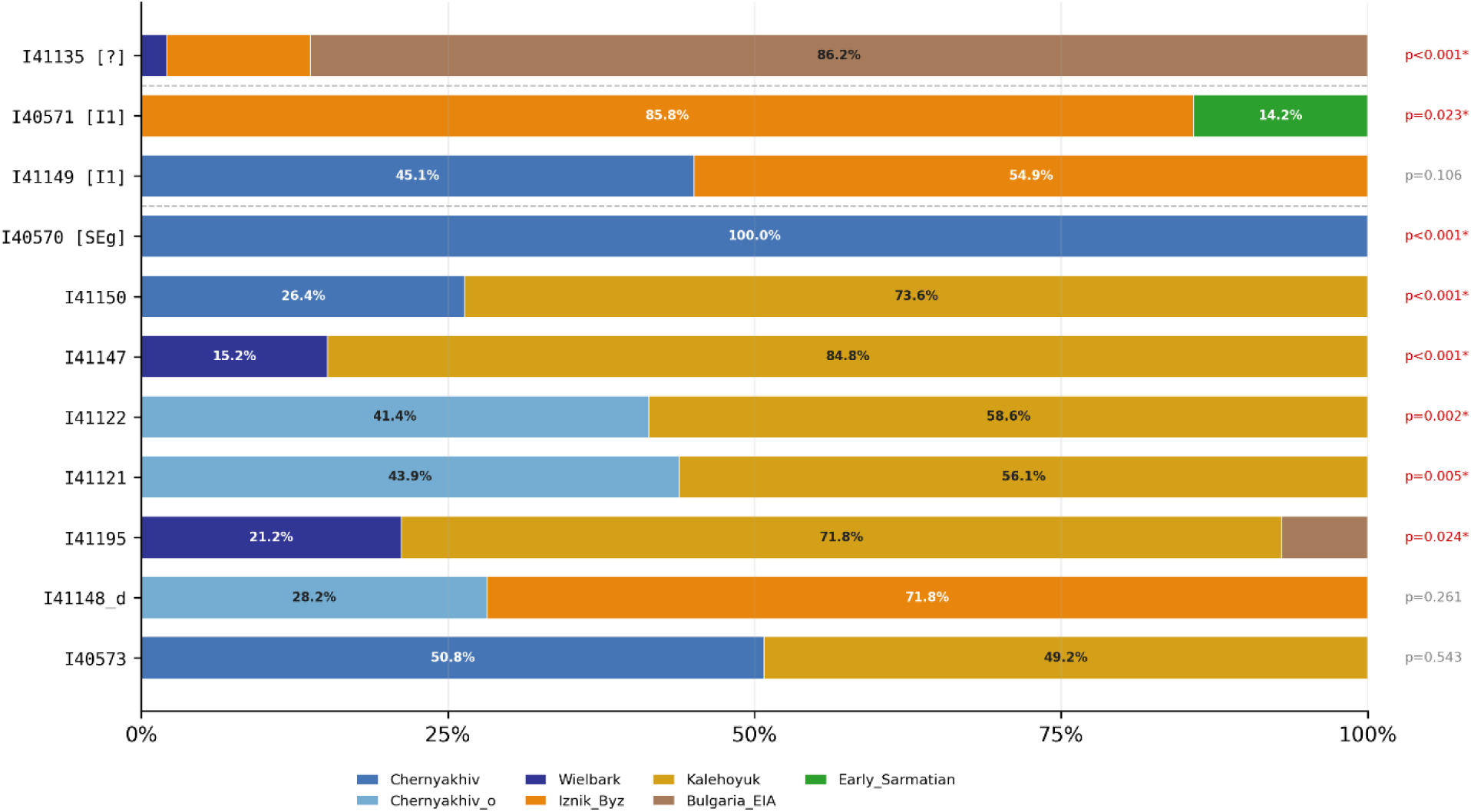
Individual-level proximal qpAdm models for the Aquae Calidae South subcluster and I1 Y-haplogroup carriers. South subcluster shows predominantly Kalehöyük-derived ancestry; I1 carriers display mixed Chernyakhov + Iznik_Byzantine profiles.

### 2.3 Admixture Timing Places North - South Mixing in the 1st Century CE

We applied DATES^14^ to estimate the timing of admixture between northern European and southern (Balkan/Anatolian) ancestry components (see Methods). Reference panels: northern sources - Pol3 (Pruszcz Gdański + Masłomęcz + Kowalewko Wielbark-associated sites; n=68) or Chernyakhov (n=3); southern sources - BalkanCore (Bulgaria_EIA, Greek IA/LBA, North Macedonian IA; n=44), AnatoliaMax (Aegean, Central, and Eastern Anatolian populations; n=37), or Mugla3 (Roman/Byzantine-period Muğla region; n=23). Targets: KalideN (n=12), AKO (n=11), pooled KalideN+AKO (n=23). Full sensitivity analysis is in Supplementary Note S3 and Table S5.

Across all well-fitting models with Wielbark northern sources, admixture dates converge on 11 - 13 generations before burial (Z-scores 4.39 - 5.26). The best model (KalideN+AKO, Pol3 northern, AnatoliaMax southern) yields 12.44±2.36 generations (Z=5.26; nrmsd=0.186; Table 4). For burials dated ∼360 CE (KalideN) and ∼460 CE (AKO), this places admixture in the 1st century BCE to early 2nd century CE - predating the earliest documented Gothic - Roman contacts (∼170 CE).

**Table 4.** DATES admixture dating results. Top models ranked by Z-score. Generation time = 29 years. nrmsd: normalized root mean squared deviation (model fit quality; values <0.5 indicate acceptable fit).

| Target | North | South | Mean (gen) | SE | Z | nrmsd | Point est. | 95% CI |
| --- | --- | --- | --- | --- | --- | --- | --- | --- |
| KalideN+AKO (n=23) | Pol3 (n=68) | AnatoliaMax (n=37) | 12.44 | 2.36 | **5.26** | 0.186 | ~49 CE | 85 BCE - 183 CE |
| KalideN (n=12) | Pol3 (n=68) | Mugla3 (n=23) | 12.74 | 2.70 | **4.73** | 0.191 | ~4 CE | 136 BCE - 144 CE |

<sup>14</sup> Narasimhan VM, Patterson N, Moorjani P, Rohland N, Bernardos R, Mallick S, et al. (2019). The formation of human populations in South and Central Asia. *Science* 365:eaat7487. doi: 10.1126/science.aat7487.

| Target | North | South | Mean (gen) | SE | Z | nrmsd | Point est. | 95% CI |
| --- | --- | --- | --- | --- | --- | --- | --- | --- |
| KalideN (n=12) | Pol3 (n=68) | BalkanCore (n=44) | 12.09 | 2.61 | **4.63** | 0.318 | ~41 CE | 97 BCE - 179 CE |
| KalideN (n=12) | Pol3 (n=68) | AnatoliaMax (n=37) | 11.01 | 2.42 | **4.55** | 0.242 | ~41 CE | 97 BCE - 179 CE |
| AKO (n=11) | Chernyakhov (n=3) | Mugla_Stratonikeia (n=10) | 17.23 | 3.94 | **4.38** | 0.454 | ~40 BCE | 294 BCE - 214 CE |
| KalideN+AKO (n=23) | Pol3 (n=68) | BalkanCore (n=44) | 12.12 | 2.76 | **4.39** | 0.221 | ~49 CE | 85 BCE - 183 CE |

Adding Scandinavian/Danish Iron Age samples to the northern panel shifts dates older by 3 - 5 generations and inflates standard errors (Pol3+Scandinavia vs. BalkanCore: 14.63±4.59 gen, Z=3.19, versus Pol3 alone: 12.09±2.61 gen, Z=4.63). Adding Polish Iron Age Przeworsk culture similarly degrades inference. The ∼12-generation admixture signal is best recovered with Wielbark-like northern proxies and degrades when broader Scandinavian Iron Age panels are substituted or added, though it does not exclude the possibility that other Wielbark-frequency-matched sources could produce comparable fits.

BalkanCore (12.09 gen, Z=4.63), AnatoliaMax (11.01 gen, Z=4.55), and Mugla3 (12.74 gen, Z=4.73) produce timing estimates differing by less than 1.1 generations - well within the SE of ∼2.5 generations. Pooling Balkan and Anatolian sources into a single southern reference degrades inference (Z drops to ∼3, SE approximately doubles), consistent with internal structure within the substrate but not with two temporally separable admixture pulses. These results are consistent with admixture involving a Balkan - Anatolian substrate already blended prior to its encounter with Wielbark-related populations.

KalideN and AKO show different optimal northern proxy preferences (KalideN: Z=4.34 for Wielbark vs. Z=2.21 for Chernyakhov; AKO: Z=4.38 for Chernyakhov vs. Z=2.64 for Wielbark), likely reflecting the larger southern fraction in KalideN creating stronger allele-frequency contrast with Wielbark. KalideN fails as a southern proxy for AKO (Z<1), ruling out an ancestor - descendant relationship between the two groups.

Control tests on non-Gothic populations suggest signal specificity. Serbia_Viminacium_Roman_Rit (n=3)^15^ and additional Roman-period Bulgarian populations failed all DATES contrasts (Z<1.8; nrmsd>0.66). The ∼12-generation signal with Z=4.4 - 5.3 is not recovered in any non-Gothic Roman-period Balkan population tested.

**Figure 5A.**
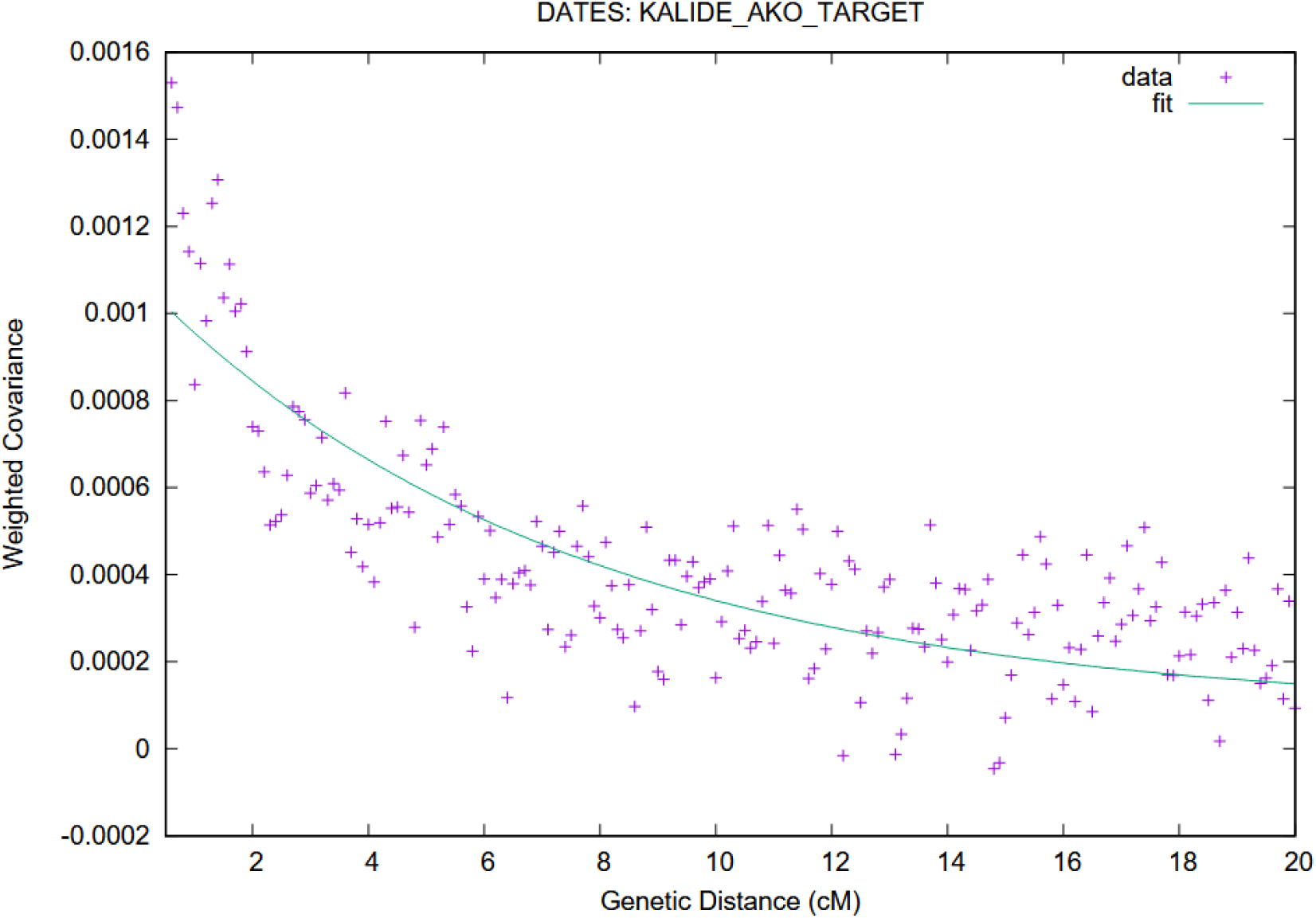
DATES weighted covariance decay curve for the best-fitting model (KalideN+AKO target; Pol3 northern source; AnatoliaMax southern source; Z=5.26, 12.44±2.36 generations). Observed decay (points) against fitted exponential (line). (B) Comparison of admixture date estimates across different southern source configurations, showing convergence at ∼12 generations:

**Figure 5B.**
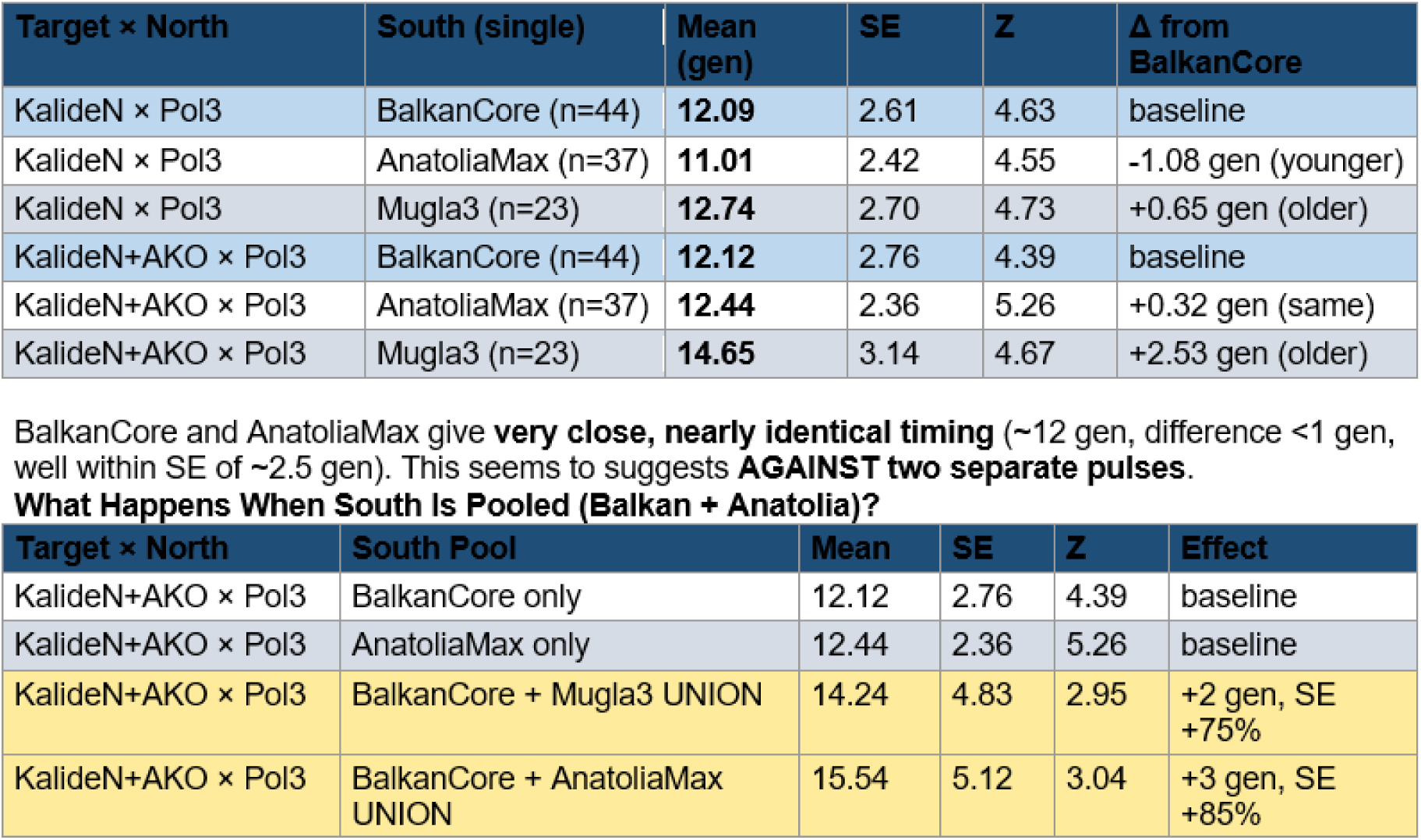
Comparison of admixture date estimates across different southern source configurations (BalkanCore, AnatoliaMax, Mugla3), showing convergence at ∼11 - 13 generations before burial across all well-fitting Wielbark-northern models.

### 2.4 Uniparental Markers Corroborate Ancestry Heterogeneity

Y-haplogroups were assigned for 19 of 24 males (79%; yHaplo, terminal SNPs ≥3× coverage, ISOGG v15.73). Only ∼23% carry Germanic-associated lineages (I1, R-DF90), below the ∼41% I1 frequency observed in Iron Age Wielbark populations ^16^ and inconsistent with direct descent from an undiluted Wielbark source. The assemblage spans multiple continental affiliations: Aquae Calidae ∼36% Anatolian/Near Eastern (J2a, G2a), ∼29% Germanic (I1, R-DF90), ∼18% Balto-Slavic (R1a), ∼12% Steppe/East Asian (C2a, Q1b). AKO shows phase-specific dynamics: C1/C2 dominated by northern lineages (∼50% I1/I2, ∼30% R1a); C4 mixed (∼40% northern, ∼30% steppe/East Asian); C5 exclusively Anatolian/Near Eastern (2/2 confirmed: G2a, J2a).

Autosomal and Y-chromosome signals are discordant in key cases. Three confirmed I1 carriers are consistent with Roman_Byzantine (28%) + Kalehöyük (26%) + Wielbark (22%) + Bulgaria_EIA (15%) + Chernyakhov (9%; p=0.695) - carrying Scandinavian-associated Y-lineages alongside substantial Anatolian autosomal ancestry. Y-haplogroup lineage does not predict autosomal ancestry composition in these assemblages. Full haplogroup assignments are in Supplementary Table S7.

**Figure 6.**
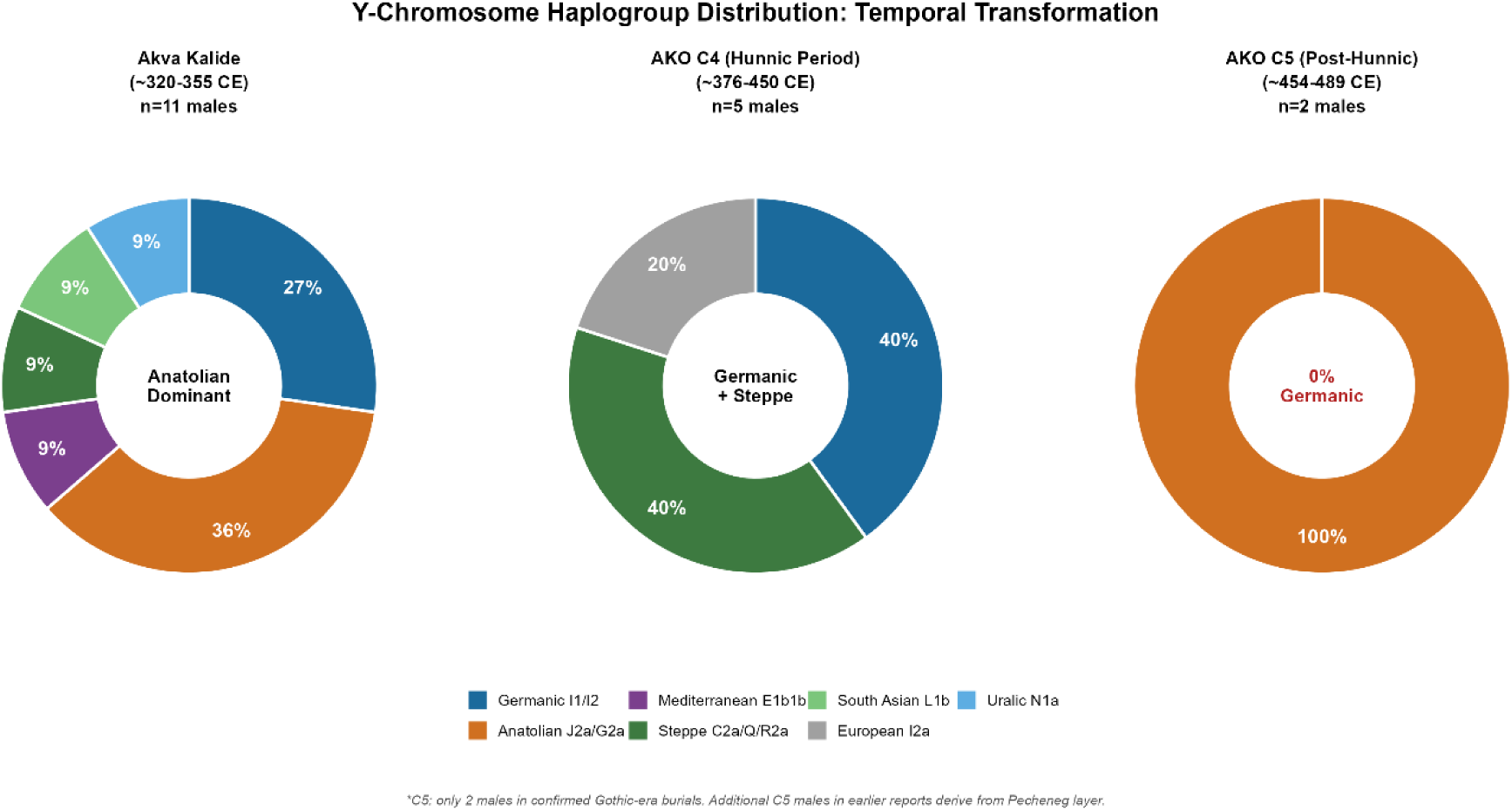
Y-chromosome haplogroup distribution across Aquae Calidae and AKO by temporal phase. Only ∼23% of the assemblage carries Germanic-associated lineages (I1, R-DF90), well below the ∼41% I1 frequency observed in Iron Age Wielbark populations^17^.

mtDNA haplogroups were assigned for all 38 individuals (HaploGrep2^18^, PhyloTree Build 17). The assemblage is predominantly local Balkan/Aegean (∼55 - 60%: H, J, T2, K, HV, V, X2, W, I), with steppe/forager (∼20 - 25%: U5, U4a2, U2e1f1), Near Eastern/Caspian (∼10 - 15%: U7b, N1a1a3, N1b1a), rare East Eurasian (C5a1; ∼3%), and African (L1b1a; ∼2%) components. Aquae Calidae displays 20 distinct mtDNA haplogroups among 20 typed individuals with no shared maternal lineages across any pair, inconsistent with a founding population. mtDNA distributions at AKO remain relatively stable across phases (∼55% Balkan throughout), contrasting with the substantial Y-chromosome phase shifts, consistent with - though not proving - male-biased demographic processes. Formal f4 sex-bias tests lack statistical power at current sample sizes.

### 2.5 Kinship Structure: Close Relatives Confined Within Sites

IBD analysis (READv2^19^; ∼220,000 pairs evaluated; cutoff 0.04) identifies 9 intra-Gothic kinship pairs spanning three degrees of relationship (3 first-degree, 3 second-degree, 3 third-degree). Eight of these pairs occur within AKO; one pair at Aquae Calidae is consistent with a third-degree relationship. No confirmed first- or second-degree kinship was detected between the two sites or between Gothic individuals and external reference populations (55 external pairs; all third-degree or more distant by confirmed degree), indicating that nuclear-family boundaries did not cross between burial communities sharing the same material and ritual repertoire. Full pairwise kinship calls and sensitivity analyses are in Supplementary Note S2 and the Supplementary IBD Workbook.

## 3. Discussion

### 3.1 Admixture Timing and the Trans-Danubian Frontier Hypothesis

Balkan and Anatolian southern sources yield indistinguishable admixture timing (∼12 generations before burial; point estimate ∼50 CE; 95% CI: 85 BCE - 183 CE), and this signal is absent from non-Gothic Roman-period Balkan controls. The temporal estimate predates the earliest documented Gothic - Roman contacts by ∼120 - 170 years, placing the admixture event earlier than the historically documented Gothic settlement south of the Danube.

One historically plausible framework consistent with the temporal constraints is admixture in a trans-Danubian frontier context, potentially including Roman Dacia, though alternative pathways remain possible. The conquest of Dacia in 106 CE created a Roman province north of the Danube that received documented Anatolian and central Balkan colonists (attested epigraphically and archaeologically^20^), generating a Balkan - Anatolian blended substrate of the type that DATES identifies as the southern component in Gothic targets. The Wielbark expansion into the Pontic region during the 2nd century CE brought populations carrying Wielbark-like ancestry into proximity with this mixed frontier population. When Rome abandoned Dacia under Aurelian (∼271 - 275 CE), both displaced frontier communities and the incoming Tervingi Gothic polity could have carried the resulting admixture signature south of the Danube.

The indistinguishable timing of Balkan and Anatolian sources is consistent with this model: if Anatolian settlers and Balkan provincials were already blended in Dacia before their encounter with Wielbark-related populations, DATES would correctly recover a single pre-mixed southern source rather than two separable pulses - which is what the pooled-source analysis suggests (Z degrading from ∼5 to ∼3 when sources are combined). We emphasize that DATES constrains the timing of admixture but cannot independently establish its geographic location; the Roman Dacia model is the scenario most consistent with the temporal evidence, but it remains a hypothesis requiring direct ancient DNA sampling from the trans-Danubian frontier zone to test.

### 3.2 Gothic Material Culture Was Practiced by Genetically Diverse Communities

The genetic heterogeneity across both sites-ancestry profiles ranging from ∼100% Chernyakhov to ∼85% Anatolian-related, with no single qpAdm model fitting the full assemblage is consistent with Gothic identity functioning as a permeable cultural-political category rather than a marker of biological descent. Three individuals illustrate the range of biological origins subsumed under shared Gothic mortuary practice. I41205 (AKO, C4) carries Y-haplogroup C2a1a (closest modern parallels among Buryats and Khalkha Mongols) with ∼6.5% East Asian autosomal ancestry. I41194 (Aquae Calidae) carries J2a Y-haplogroup alongside mtDNA L1b1a, a sub-Saharan African lineage. I40570 (Aquae Calidae) is consistent with ∼50% ancestry from Levant/Egypt-related sources, with J1 Y-haplogroup, and falls entirely outside the main site PCA cluster. All three were buried with diagnostic Gothic artifacts and Christian orientation.

The observed heterogeneity is consistent with “ethnogenesis” models of Gothic affiliation ^21,22^, but cannot by itself prove the particular institutional mechanism of identity formation or maintenance as suggested by Wolfram and Pohl (Arian Christianity, Gothic language codified by Ulfilas and foederati legal status). The absence of close kinship between the two sites reinforces this picture: nuclear-family boundaries did not cross between communities that nonetheless shared a similar material and ritual repertoire while standing apart from the surrounding Roman provincial culture. Historical scholarship offers several mechanisms by which Gothic affiliation may have been reproduced across populations of diverse ancestry, including ecclesiastical, linguistic, and political institutions; our data are compatible with such models yet cannot directly discriminate among them.

### 3.3 Temporal Dynamics at AKO

AKO’s sequential necropoleis document progressive northward-ancestry dilution: EHG declines from 27.9±3.2% (C1, ∼350 CE) to 4.6 - 16.2±4.1% (C5, ∼454 - 489 CE) as Anatolian Neolithic Farmer ancestry rises from ∼21% to ∼32 - 50%. Y-chromosome composition shifts from ∼50% Germanic-associated lineages (C1/C2) to exclusively Anatolian/Near Eastern haplogroups in C5, while mtDNA remains stable (∼55% Balkan throughout). The contrast between shifting Y-chromosome composition and comparatively stable mtDNA composition is suggestive of male-biased demographic processes, though current sample sizes do not allow a firm inference. The genomic trajectory - progressive dilution rather than abrupt replacement is more compatible with long-term local integration under Theoderic Strabo’s foederati communities (∼440s CE onward; see “Thracian Goths” in Heather^23^) than with the narrower chronological window of Theodoric the Amal (474 - 488 CE). C5 dating rests partly on Zeno coins (474 - 491 CE); without independent radiocarbon dates, the historical association remains suggestive.

### 3.4 Limitations

Phase sample sizes, particularly C5 (n=3), limit statistical power. Reference panel gaps - no published Roman-era Central Anatolian aDNA; Chernyakhov samples limited to Ukrainian sites required temporally or geographically approximate proxies. Genome coverage constrains Y-haplogroup subclade resolution and IBD sensitivity. DATES assumes pulse admixture and cannot independently determine the geographic location of the admixture event. Genetic heterogeneity consistent with ethnogenesis does not prove that model; similar patterns could arise from military conscription, serial migrations, or exogamy without shared ethnic identification. Expanded control sampling from published Roman-period Balkan datasets would further test the specificity of the Gothic-associated DATES signal.

### 3.5 Implications for Genetic Studies of Migration Period Identity

This study demonstrates both the power and the limitation of ancient DNA research as a methodological instrument in addressing questions of ethnic identity. The power is chronometric: DATES places a specific quantitative constraint on when the north - south ancestry mixture occurred, identifies it as more consistent with Wielbark-like than generic northern European ancestry, demonstrates that Balkan and Anatolian sources entered the Gothic gene pool as a pre-blended substrate rather than as separate events, and shows that this signature is absent from non-Gothic contemporaries in the same region. None of these observations was accessible to archaeology or historical linguistics alone, and together they narrow the range of historically plausible scenarios for Gothic population history. The limitation is however instructive: the data that place admixture before the first Gothic - Roman contacts cannot determine whether that admixture happened in Roman Dacia, along the Pontic frontier, or through a pathway not yet considered.

The extreme biological heterogeneity of the two assemblages cannot by itself be distinguished from the products of military conscription, commercial networks, or serial migrations that left no shared ethnic trace. We suggest that what ancient DNA can add to Migration Period studies is not an ultimate resolution of identity questions but the replacement of hard-to-verify narrative assumptions with testable chronometric and demographic constraints.

The identities themselves, the institutional frameworks through which populations recognized and reproduced “Gothic” belonging remain in the domain of the textual, epigraphic, and material record that genomics can constrain but hardly supplant.

Gothic-associated communities sampled at Aquae Calidae and AKO were not biologically coherent populations but genetically heterogeneous assemblages linked by shared mortuary practice and historical affiliation. Their distinct ancestry profiles, together with admixture dates centering ∼11 - 13 generations before burial, indicate that the northern and southern components of these groups were brought together before the historically documented Gothic settlement south of the Danube. These findings narrow the range of plausible demographic scenarios for Gothic population history in the Balkans while also underscoring a central limitation of paleogenomics as a method per se: ancient DNA can constrain when and between which ancestry streams mixture occurred, but it cannot by itself determine the institutional and social processes through which “Gothic” belonging was created, maintained, and recognized.

## 4. Methods

### Sample Collection

Bone and dental samples from 38 individuals (Aquae Calidae: n=23; AKO: n=15) were collected from archaeological excavations conducted by the National Museum of History, Sofia, and the Regional Historical Museum, Burgas.

### Ancient DNA Data Generation

Wet laboratory work was performed in the ancient DNA laboratory at Harvard Medical School in Boston, USA. DNA extraction followed Rohland et al.^24^, using silica beads with Dabney buffer in a robotic workflow. Libraries were prepared using two protocols depending on sample quality: double-stranded libraries (n=74 libraries from 37 individuals) with partial UDG treatment following Rohland et al.^25^, and single-stranded libraries (n=4 libraries from 2 individuals) with USER treatment following Gansauge et al.^26^. All libraries were enriched by in-solution capture targeting the Twist Ancient DNA panel using the Rohland et al.^27^ reagents (Twist1.44Mv1.5 for double-stranded libraries; Twist1.4Mv1.8 for single-stranded libraries). Double-stranded libraries were sequenced on an Illumina NovaSeq X 10B instrument; single-stranded libraries were sequenced on an Illumina NovaSeq X 25B instrument. Library-level quality metrics, pass/fail assessments, and per-sample coverage statistics are reported in Supplementary Table S7.

### Ancient DNA Bioinformatic Processing

The great majority of data come from sequencing the products of in-solution enrichment targeting more than one million known polymorphisms. Libraries were marked with identification tags (barcodes and indices) before sequencing in pools. We merged paired-end sequences, requiring no more than one mismatch in the overlap between paired sequences where base quality is at least 20, or three mismatches if base quality is <20; sequences that could not be merged were not analyzed. Adapters and identification tags were stripped using a custom toolkit (ADNA-Tools v2.1.0). Merged sequences were aligned to the hg19 human reference genome with decoy sequences (hs37d5) using BWA SAMSE v0.7.15 with parameters -n 0.01 -o 2 and -l 16500. Duplicate reads were marked using Picard MarkDuplicates v2.17.10. Merged sequences were additionally mapped to the mitochondrial DNA Reconstructed Sapiens Reference Sequence (RSRS) for mitochondrial-specific metrics. Bioinformatic processing produced key quality metrics including authenticity estimates based on elevated damage rates at sequence ends, contamination rates, and endogenous content rates. Pseudo-haploid genotypes were called by randomly sampling one allele per site.

### Principal Component Analysis

PCA was conducted using smartpca (EIGENSOFT v7.2.1)^28^, projecting ancient samples onto principal components computed from 1,240 modern West Eurasian individuals (shrinkmode: YES; lsqproject: YES). PC1 and PC2 captured approximately 19.9% and 9.9% of total variance.

### f3 and f4 Statistics

Outgroup f3 statistics [f3(Mbuti; Target, Reference)] and f4 statistics^29^ were computed using admixtools2^30^. Standard errors were estimated by jackknife over genomic blocks. Full results are in Supplementary Tables S1 and S6 as f3 results are additionally discussed in Supplementary Note S5.

### qpAdm Ancestry Modeling

qpAdm modeling was implemented in admixtools2. Right populations: Mbuti.DG, Israel_Natufian.SG, Russia_Yana_UP.SG, Russia_MA1_HG.SG, Russia_Kostenki14.SG, Italy_North_Villabruna_HG.SG, Morocco_Iberomaurusian.SG, Turkey_Boncuklu_PPN.SG, Iran_GanjDareh_N.SG, Karitiana.DG, Russia_DevilsCave_N.SG. Models with p > 0.05 were considered feasible. Source rotation testing encompassed ∼100 ancient Anatolian samples from >20 populations spanning Roman, Byzantine, Iron Age, and Bronze Age periods. Full model outputs are in Supplementary Tables S2 - S4.

### DATES Admixture Dating

DATES^31^ was applied with parameters: binsize=0.001, mindis=0.02, maxdis=0.5, qbin=10, jackknife=YES, seed=77, runfit=YES, afffit=YES, lovalfit=0.6, minparentcount=2, numchrom=22. Generation time was set to 29 years. Full sensitivity analysis, reference panel definitions, and control population results are in Supplementary Note S3 and Table S5.

### Uniparental Markers

Y-haplogroups were assigned using yHaplo (terminal SNPs with ≥3× coverage; ISOGG v15.73).

mtDNA haplogroups were assigned using HaploGrep2^32^ (PhyloTree Build 17).

### IBD and Kinship Analysis

IBD analysis was performed using READv2^33^ on ∼220,000 pairs with a kinship cutoff of 0.04. Full pairwise kinship calls and sensitivity analyses are in Supplementary Note S2 and the Supplementary IBD Workbook.

## Supporting information

Supplemental Table S6 F4 statistics

Supplemental Table S7 Sample Information

Supplemental IBD Workbook

Supplemental Note S2 IBD

Supplemental Note S3 DATES

Supplemental Note S4 Archaeology Timeline

Supplemental Note S5 F3 Statistics

## Data Availability

Raw sequencing data: European Nucleotide Archive (PRJXXXXXX). Genotypes: Allen Ancient DNA Resource (AADR). Interactive PCA, f3, and IBD visualizations: [GitHub repository URL].

## Acknowledgments

This publication has been developed within the framework of project BG05SFPR001-3.004-0005-C01 “Achieving Innovation and Growth through Doctoral Research”, implemented under Programme “Education” 2021–2027, co-financed by the European Union through the European Social Fund Plus (ESF+).

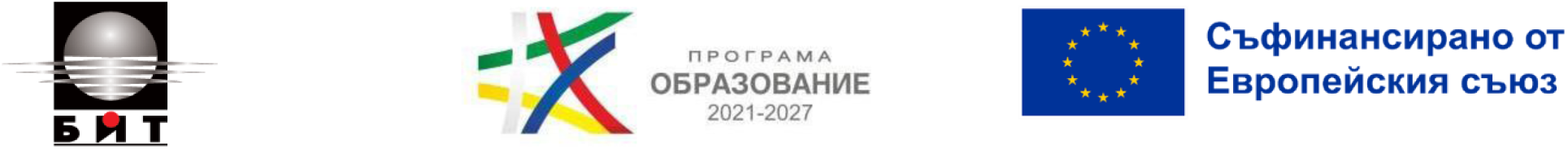

This research was conducted as part of the “Epochs of History” paleogenomic project.

This work was supported by the Bulgarian National Science Fund (grant/contract no. KP-06-N70/8; Contract № КП-06-H70/8) as an interdisciplinary collaboration between the IBCT - Bulgarian Academy of Science, UNIBIT – Sofia, the National Institute for Archaeology with Museum, National Historical Museum, Bulgarian Regional Museums of History and Harvard University’s David Reich Laboratory of Archaeogenetics.

We are deeply grateful to Prof. Stoyan Denchev (UNIBIT, Sofia) for his unwavering support as doctoral adviser to SS and for generously providing access to the computational server infrastructure on which many of the analyses in this study were performed.

We extend our heartfelt thanks to Prof. Boni Petrunova (National Museum of History, Sofia) for hosting the research at National Museum of History, for facilitating access to archaeological collections, and for her sustained support and encouragement throughout the project. This work would not have been possible without her institutional vision and personal commitment.

We thank Prof. Rumyana Preshlenova (Institute for Balkan Studies with Centre for Thracology, Bulgarian Academy of Sciences / IBCT-BAS) for her support of the interdisciplinary framework within which this research was conducted.

We gratefully acknowledge Aisling Kearns and the ancient DNA clean-room staff at the Reich Laboratory, Harvard Medical School, for expert sample processing and library preparation, and the Harvard sequencing facility for high-throughput sequencing support.

## Author Contributions

S.S.: Conceptualization, Formal analysis, Data curation, Methodology, Software (bioinformatics pipeline implementation), Visualization, Writing – original draft, Writing – review & editing, Project administration. T.C.: Conceptualization, Funding acquisition, Resources (archaeological and chronological framework), Writing – original draft (Supplementary Note S4), Writing – review & editing. T.W.: Formal analysis, Methodology, Writing – review & editing. K.S.: Resources (archaeological collections, Veliki Preslav assemblage), Investigation. D.M.: Resources (archaeological collections, Regional Historical Museum Burgas), Investigation. D.M.: Resources (archaeological collections, Regional Historical Museum Burgas), Investigation. M.K.: Resources (National Archaeological Institute collections and site documentation), Investigation. G.S.: Writing. M.N.: Resources (Regional Historical Museum Burgas collections), Investigation. D.N.: Investigation (skeletal sampling and physical specimen collection), Resources. P.H.: Writing – review & editing (historical interpretation and ethnogenesis framework). D.I.T.: Writing – review & editing. M.Z.: Writing – review & editing. I.L.: Methodology, Formal analysis, Writing – review & editing, Supervision. D.R.: Conceptualization, Funding acquisition, Resources (sequencing and laboratory infrastructure), Supervision, Writing – review & editing.

## Footnotes

1 Heather P (1996). *The Goths*. Blackwell, Oxford. ISBN 9780631165361.

2 Stolarek I, Handschuh L, Juras A, Nowaczewska W, Piontek J, Kozlowski P, Figlerowicz M (2023). Genetic history of East-Central Europe in the first millennium CE. *Genome Biology* 24:173. doi: 10.1186/s13059-023-03013-9.

3 Saag L, Utevska O, Zadnikov S, Shramko I, Gorbenko K, Bandrivskyi M, et al. (2025). North Pontic crossroads: Mobility in Ukraine from the Bronze Age to the early modern period. *Science Advances* 11(2):eadr0695. doi: 10.1126/sciadv.adr0695.

4 Jordanes (6th century CE). Getica (The Origin and Deeds of the Goths). Trans. C. C. Mierow (1915). Princeton University Press, Princeton.

5 Heather P (1996). *The Goths*. Blackwell, Oxford. ISBN 9780631165361.

6 Wolfram H (1988). *History of the Goths*. University of California Press, Berkeley. ISBN 9780520052598.

7 Pohl W, Heydemann G, eds. (2013). *Strategies of Identification: Ethnicity and Religion in Early Medieval Europe*. Brepols, Turnhout. ISBN 9782503533841.

8 Veeramah KR, Scheib CL, Kirsanow K, Sell C, Alber C, Flaig J, et al. (2018). Population genomic analysis of elongated skulls reveals extensive female-biased immigration in Early Medieval Bavaria. *Proceedings of the National Academy of Sciences* 115(13):3494–3499. doi: 10.1073/pnas.1719880115.

9 Jordanes (6th century CE). Getica (The Origin and Deeds of the Goths). Trans. C. C. Mierow (1915). Princeton University Press, Princeton.

10 Procopius (6th century CE). On Buildings (De Aedificiis). Trans. H. B. Dewing, G. Downey. Loeb Classical Library 343. Harvard University Press, Cambridge MA. doi: 10.4159/DLCL.procopius-buildings.1940.

11 Thompson EA (2008). *The Visigoths in the Time of Ulfila*. 2nd edn. Duckworth, London. ISBN 9780715637005.

12 Heather P (1986). The crossing of the Danube and the Gothic conversion. *Greek, Roman and Byzantine Studies* 27(3):289–318.

13 Kulikowski M (2007). *Rome’s Gothic Wars: From the Third Century to Alaric*. Cambridge University Press, Cambridge. ISBN 9780521846332.

14 Narasimhan VM, Patterson N, Moorjani P, Rohland N, Bernardos R, Mallick S, et al. (2019). The formation of human populations in South and Central Asia. *Science* 365:eaat7487. doi: 10.1126/science.aat7487.

15 Olalde I, Carrion P, Mikic I, Rohrlach AB, Mallick S, Lazaridis I, et al. (2023). A genetic history of the Balkans from Roman frontier to Slavic migrations. *Cell* 186(25):5472–5485.e9. doi: 10.1016/j.cell.2023.10.018.

16 Stolarek I, Handschuh L, Juras A, Nowaczewska W, Piontek J, Kozlowski P, Figlerowicz M (2023). Genetic history of East-Central Europe in the first millennium CE. *Genome Biology* 24:173. doi: 10.1186/s13059-023-03013-9.

17 Stolarek I, Handschuh L, Juras A, Nowaczewska W, Piontek J, Kozlowski P, Figlerowicz M (2023). Genetic history of East-Central Europe in the first millennium CE. *Genome Biology* 24:173. doi: 10.1186/s13059-023-03013-9.

18 Weissensteiner H, Pacher D, Kloss-Brandstatter A, Forer L, Specht G, Bandelt HJ, Kronenberg F, Salas A, Schonherr S (2016). HaploGrep 2: mitochondrial haplogroup classification in the era of high-throughput sequencing. *Nucleic Acids Research* 44(W1):W58–W63. doi: 10.1093/nar/gkw233.

19 Monroy Kuhn JM, Jakobsson M, Gunther T (2018). Estimating genetic kin relationships in prehistoric populations. *PLOS ONE* 13(4):e0195491. doi: 10.1371/journal.pone.0195491.

20 Oltean IA (2007). Dacia: Landscape, Colonisation, Romanisation. Routledge Monographs in Classical Studies. Routledge, London/New York. ISBN 9780415412520.

21 Wolfram H (1988). *History of the Goths*. University of California Press, Berkeley. ISBN 9780520052598.

22 Pohl W, Heydemann G, eds. (2013). *Strategies of Identification: Ethnicity and Religion in Early Medieval Europe*. Brepols, Turnhout. ISBN 9782503533841.

23 Heather P (1996). *The Goths*. Blackwell, Oxford. ISBN 9780631165361.

24 Rohland N, Glocke I, Aximu-Petri A, Meyer M (2018). Extraction of highly degraded DNA from ancient bones, teeth and sediment for high-throughput sequencing. *Nature Protocols* 13(11):2447–2461. doi: 10.1038/s41596-018-0050-5.

25 Rohland N, Harney E, Mallick S, Nordenfelt S, Reich D (2015). Partial uracil-DNA-glycosylase treatment for screening of ancient DNA. *Philosophical Transactions of the Royal Society B* 370:20130624. doi: 10.1098/rstb.2013.0624.

26 Gansauge M-T, Aximu-Petri A, Nagel S, Meyer M (2020). Manual and automated preparation of single-stranded DNA libraries for the sequencing of DNA from ancient biological remains and other sources of highly degraded DNA. *Nature Protocols* 15(8):2279–2300. doi: 10.1038/s41596-020-0338-0.

27 Rohland N, Mallick S, Mah M, Maier R, Patterson N, Reich D (2022). Three assays for in-solution enrichment of ancient human DNA at more than a million SNPs. *Genome Research* 32(11–12):2068–2078. doi: 10.1101/gr.276728.122.

28 Patterson N, Price AL, Reich D (2006). Population structure and eigenanalysis. *PLOS Genetics* 2(12):e190. doi: 10.1371/journal.pgen.0020190.

29 Patterson N, Moorjani P, Luo Y, Mallick S, Rohland N, Zhan Y, Genschoreck T, Webster T, Reich D (2012). Ancient admixture in human history. *Genetics* 192(3):1065–1093. doi: 10.1534/genetics.112.145037.

30 Maier R, Flegontov P, Flegontova O, Isildak U, Changmai P, Reich D (2023). On the limits of fitting complex models of population history to f-statistics. *eLife* 12:e85492. doi: 10.7554/eLife.85492. Software: admixtools2, https://github.com/uqrmaie1/admixtools.

31 Narasimhan VM, Patterson N, Moorjani P, Rohland N, Bernardos R, Mallick S, et al. (2019). The formation of human populations in South and Central Asia. *Science* 365:eaat7487. doi: 10.1126/science.aat7487.

32 Weissensteiner H, Pacher D, Kloss-Brandstatter A, Forer L, Specht G, Bandelt HJ, Kronenberg F, Salas A, Schonherr S (2016). HaploGrep 2: mitochondrial haplogroup classification in the era of high-throughput sequencing. *Nucleic Acids Research* 44(W1):W58–W63. doi: 10.1093/nar/gkw233.

33 Monroy Kuhn JM, Jakobsson M, Gunther T (2018). Estimating genetic kin relationships in prehistoric populations. *PLOS ONE* 13(4):e0195491. doi: 10.1371/journal.pone.0195491.

