## Supplemental Note S2 IBD for "Paleogenomic Evidence for Genetic Heterogeneity and Prior Admixture in Gothic-Associated Communities of Late Antique Bulgaria"

Supplementary Note S2

**Kinship and allele-sharing analysis of Gothic-associated populations in Late Antique Bulgaria**

This supplementary note presents detailed READv2-based kinship and allele-sharing results that complement the manuscript sections on kinship (§2.5) and the broader discussion of population heterogeneity and Gothic affiliation (§3.2, §3.5). Throughout this note, “IBD/kinship” is used as shorthand for allele-sharing-based relationship inference; the primary estimates reported here are generated with READv2 rather than phased haplotype tract detection. The results are most informative about within-site kinship clustering and the absence of confirmed close external ties in the current sample. Population-enrichment and temporal-distribution analyses are retained as exploratory summaries of relative affinity and should be read as hypothesis-generating rather than as direct evidence for specific historical scenarios.

### S2.1 Analytical framework

Pairwise kinship coefficients were estimated for all individuals using READv2 (Monroy Kuhn et al. 2018), which infers biological relatedness from patterns of allele sharing without requiring phased haplotypes. The analysis encompassed approximately 220,000 pairwise comparisons involving Gothic-associated individuals from two primary sites: Aquae Calidae (Akva Kalide, southeastern Bulgaria) and the Aul of Khan Omurtag (AKO, northeastern Bulgaria).

We used a kinship coefficient threshold of θ ≥ 0.04, which is broadly compatible with third-degree and more distant relationships but may also reflect background population structure at the lower end of the range. Samples with fewer than 15,000 overlapping SNPs were excluded from the main analysis. Low-confidence classifications were flagged where SNP overlap was insufficient for reliable degree assignment. First-degree relationships (θ ≈ 0.125–0.25) correspond to parent-offspring or full siblings; second-degree relationships (θ ≈ 0.0625–0.125) to grandparent-grandchild, half-siblings, or avuncular pairs; third-degree relationships (θ ≈ 0.04–0.0625) to first cousins or comparable genealogical distance.

Population-size normalized enrichment scores were calculated as the ratio of observed connections to expected connections based on each population’s representation in the total comparison dataset (n = 979 individuals from merged .fam files). Enrichment values are useful for comparing relative overrepresentation across reference groups, but they are sensitive to grouping strategy, sample size, and the composition of the reference panel. Individuals with more than five Gothic connections were removed in sensitivity analyses to test the effect of highly connected samples.

### S2.2 Internal Gothic kinship structure

Within the Gothic-associated sample, nine kinship pairs were identified across the two Bulgarian sites (Table S2-1; Supplementary IBD Workbook: Internal Gothic Kinship sheet). Eight of these pairs occur within AKO, whereas Aquae Calidae contains a single third-degree pair. This asymmetry is robust in the present dataset and indicates that the sampled AKO burials include a denser representation of close relatives than the sampled Aquae Calidae burials. However, the difference should not be over-interpreted: it may reflect variation in burial selection, cemetery use, sampling density, or community structure, and it does not by itself determine a unique settlement model.

| **Relationship degree** | **Total** | **AKO** | **Aquae Calidae** |
| --- | --- | --- | --- |
| First degree | 3 | 3 | 0 |
| Second degree | 3 | 3 | 0 |
| Third degree | 3 | 2 | 1 |
| Total | 9 | 8 (89%) | 1 (11%) |

*Table S2-1. Internal kinship pairs by site.*

At AKO, the presence of first-, second-, and third-degree relationships within the cemetery is consistent with repeated use of the burial ground by related individuals over more than one generation. This result supports the manuscript’s inference that some AKO burials sampled closely connected family groups within the site.

At Aquae Calidae, the near-absence of close internal kinship in the sampled individuals is compatible with a more dispersed burial catchment, a more selective sample of burials, or lower representation of close relatives in the sequenced subset. In the current dataset, it is safer to describe Aquae Calidae as kinship-sparse rather than to infer a specific social or institutional model from this pattern alone.

### S2.3 Absence of confirmed close external relationships in the current sample

No confirmed first-degree or high-confidence second-degree relationships were detected between Gothic-associated individuals and non-Gothic reference individuals (Table S2-2; Supplementary IBD Workbook: All Gothic Pairs sheet). The external connections that were identified are overwhelmingly third-degree or more distant. In the present dataset, this pattern indicates that the sampled burial communities are not linked to the available external comparators by close documented family ties.

| **Relationship type** | **Gothic-other pairs** | **1st deg.** | **2nd deg.** | **3rd deg. or more distant** |
| --- | --- | --- | --- | --- |
| First degree | 0 | 0 | — | — |
| Second degree (HC) | 0 | — | 0 | — |
| Second degree (LC) | 4 | — | 4 | — |
| Third degree | 25 | — | — | 25 |
| 4th degree+ | 26 | — | — | 26 |

*Table S2-2. External kinship calls involving Gothic-associated individuals.*

This pattern is compatible with an absence of close family ties between the sampled Gothic-associated burial groups and the currently available external reference individuals. At the same time, it should not be treated as direct proof of strict endogamy or of any specific institutional mechanism of social boundary maintenance. Small community size, incomplete reference coverage, and sampling effects could also contribute to the absence of close external pairs.

Sensitivity analyses did not change the central result. Relaxing the kinship threshold to θ ≥ 0.02 increased the number of distant pairs but did not produce new confirmed first- or second-degree external ties. Tightening the threshold to θ ≥ 0.06 reduced the number of calls overall while preserving the absence of close external relationships. Re-analysis of the higher-coverage subset likewise retained internal kinship clustering without generating new close external pairs.

### S2.4 Summary of external affinity signals

Population-normalized enrichment scores and temporal-distribution summaries are provided in the Supplementary IBD Workbook (Population Enrichment, Network Statistics, and Temporal Distribution sheets) and are not reproduced in full here. In brief, both sites show relative overrepresentation of connections to Pontic steppe-related, Scandinavian-related, and Balkan Antiquity reference groups, a pattern consistent with the ancestry inferences from qpAdm and DATES in the main text. Enrichment values for small reference groups (n < 15) are unreliable and should be interpreted with caution. The temporal distribution of external connections shows comparatively fewer calls in the 1st–3rd centuries CE than in adjacent temporal categories; this observation may partly reflect uneven reference panel coverage and is treated as exploratory background rather than a stand-alone finding. Full tabular outputs are available in the Supplementary IBD Workbook.

### S2.5 Site-level differences between Aquae Calidae and AKO

The kinship analyses suggest that the two Bulgarian sites differ in how close relatives are represented within the sequenced sample (Supplementary IBD Workbook: Site-Specific Comparison sheet). AKO contains several close-kin pairs across multiple relationship degrees, whereas Aquae Calidae does not. This contrast supports the manuscript’s conclusion that the two burial assemblages are socially and biologically distinct, but it should not be reduced to a single deterministic model of settlement or community organization.

| **Characteristic** | **Aquae Calidae** | **AKO** | **Reviewer-safe interpretation** |
| --- | --- | --- | --- |
| Close internal kinship | 1 third-degree pair | 8 pairs across 1st–3rd degree | AKO includes denser sampling of close relatives |
| External affinity profile | Broad external profile; strong Pontic and southern signals | Broad external profile; strong Pontic and southern signals | Both sites share wider connectivity but differ in internal clustering |
| Roman/medieval connection counts | Exploratory signals present | Exploratory signals present | Useful for comparison, but highly sensitive to panel composition |
| Working summary | Kinship-sparse sampled burial set | Kinship-clustered sampled burial set | Contrast is real; specific settlement scenarios remain open |

*Table S2-5. Site-level comparison of internal kinship and external-affinity patterns.*

For Aquae Calidae, the present sample is best described as kinship-sparse. This may reflect a dispersed burial catchment, selective excavation or sequencing, or a cemetery used by individuals drawn from multiple backgrounds. The current data do not allow these alternatives to be distinguished confidently.

For AKO, the repeated occurrence of close relatives indicates that the sampled burials include family-linked individuals across more than one generation. This supports the view that the cemetery captured a locally structured community, but it does not by itself establish whether that structure derived from continuous residence, clustered burial of certain lineages, or other site-specific mortuary practices.

### S2.6 Integration with autosomal and uniparental evidence

First, the kinship results are compatible with the manuscript’s inference that a substantial part of the southern Balkan/Anatolian-related ancestry was acquired before settlement at the Bulgarian sites. The absence of confirmed close external ties does not exclude local gene flow, but it is more compatible with a model in which much of the major ancestry mixture predates the final burial contexts sampled here.

Second, the external-affinity profiles are consistent with the autosomal and uniparental heterogeneity documented in the main text. The Gothic-associated assemblage does not behave as a biologically uniform population; rather, different individuals and subgroups combine northern, southern, and steppe-related signals to varying degrees.

Third, the kinship data reinforce the manuscript’s central interpretive point without overextending it. Shared Gothic-associated mortuary practice does not correspond to a single ancestry profile, and the lack of close kinship links between Aquae Calidae and AKO shows that culturally similar burial communities need not form one extended biological network. These results are consistent with Gothic affiliation functioning as a cultural-political framework spanning populations of diverse ancestry, while remaining insufficient to identify the precise ecclesiastical, linguistic, legal, or social mechanisms through which such affiliation was reproduced.

### S2.7 Limitations

Several limitations qualify these findings. First, READv2 estimates relatedness from allele sharing without phased haplotypes, which reduces precision for distant relationships and means that some lower-degree calls may reflect background structure rather than recent genealogy. Low-confidence classifications were flagged, but false positives and false negatives remain possible.

Second, the enrichment and temporal-distribution analyses are sensitive to the composition and grouping of the reference panel. Alternative grouping schemes or expanded temporal coverage, especially for Roman-period Balkan populations, could modify specific values and attenuate apparent gaps.

Third, the geographic and chronological interpretation of external affinities remains indirect. Enrichment values summarize relative overrepresentation, not migration routes, settlement types, or identity categories. These results should therefore be treated as complementary to the autosomal and DATES analyses, not as stand-alone historical proof.

Fourth, while the absence of confirmed close external ties is compatible with limited close intermarriage in the current sample, it cannot by itself establish endogamy. Small sample sizes, cemetery-specific structure, and incomplete reference coverage remain viable alternative explanations.

Finally, the normalization denominator (n = 979) reflects the merged comparison dataset used here; absolute enrichment values would change if the comparison set were updated, although the major qualitative contrasts are likely to be more stable than the exact numerical multipliers.
