## Supplemental Note S3 DATES for "Paleogenomic Evidence for Genetic Heterogeneity and Prior Admixture in Gothic-Associated Communities of Late Antique Bulgaria"

**Supplementary Note S3**

*DATES Admixture Dating: Sensitivity Analysis and Model Testing*

### S3.1 Overview and Analytical Framework

This note documents the complete set of DATES admixture dating analyses performed for the Gothic-associated populations from Aquae Calidae and the Aul of Khan Omurtag (AKO). The main text (§2.3) reports the principal findings; this supplement provides the full sensitivity analysis, the complete set of tested configurations including failed and diagnostic runs, formal pool definitions, and the detailed evidence underlying the pre-mixed substrate model. The summary results table is provided in Supplementary Table S5.

The DATES method (Narasimhan et al. 2019) infers admixture timing from the decay of weighted ancestry covariance along the genome. It requires specification of two reference populations representing the mixing sources and a target admixed population. The method does not require phased genomes and is applicable to low-coverage ancient DNA data; its sensitivity depends primarily on the sample size of the target population and the allele frequency contrast between the two reference sources. All runs used consistent parameters: binsize=0.001, mindis=0.02, maxdis=0.5, qbin=10, jackknife=YES, seed=77, runfit=YES, afffit=YES, lovalfit=0.6, minparentcount=2, numchrom=22.

The analyses addressed three questions posed during internal review: (i) whether pooling targets and sources could reduce standard errors without biasing estimates; (ii) whether the inferred admixture date remained stable under sensible proxy substitutions; and (iii) whether the detected signal reflected specifically Wielbark-associated ancestry or generic northern European Iron Age ancestry. A fourth question, whether the Balkan and Anatolian components of the southern substrate entered the Gothic gene pool as a single pre-mixed population or as temporally separable pulses emerged from the sensitivity results and is treated in detail in §S3.4.

### S3.2 Reference Panel Construction

#### *S3.2.1 Target populations*

Three target configurations were tested. KalideN (n=12) comprises the northern-shifted subcluster of Aquae Calidae individuals, identified by PCA position and qpAdm ancestry proportions (§2.1–2.2 of the main text). AKO (n=11) comprises all Gothic-associated individuals from the Aul of Khan Omurtag with sufficient coverage for DATES analysis. The pooled target KalideN+AKO (n=23) combines both sites to maximize statistical power. Additionally, KalideS (n=5), the southern-shifted subcluster at Aquae Calidae, was tested separately as a heterogeneity control.

#### *S3.2.2 Northern source pools*

| **Pool name** | **Composition** | **n** | **Rationale** |
| --- | --- | --- | --- |
| Pol3 (Wielbark) | Pruszcz Gdański + Masłomęcz + Kowalewko (Wielbark IA) | 68 | Core Wielbark-associated sites; maximizes contrast with southern sources |
| Pol3+Scandinavia | Pol3 + Denmark_IA + Sweden_IA + Sweden_south_PreViking + Gotland_PreViking | 82 | Tests whether signal is generic northern European or Wielbark-specific |
| Pol3+Denmark | Pol3 + Denmark_IA | 72 | Decomposition: Denmark contribution |
| Pol3+Sweden/Gotland | Pol3 + Sweden_IA + Sweden_south_ PreViking + Gotland_PreViking | 78 | Decomposition: Sweden/Gotland contribution |
| Scandinavia only | Denmark_IA + Sweden_IA + Sweden_south_PreViking + Gotland_PreViking | 14 | Tests Scandinavian-only contrast (no Wielbark) |
| Chernyakhov | Ukraine_MigrationPeriod_ Chernyakhiv (main + outlier) | 3 | Already partially admixed; tests intermediate proxy |
| Kowalewko (single site) | Poland_Kowalewko_Wielbark_IA | var. | Single-site baseline for pooling comparison |

#### *S3.2.3 Southern source pools*

| **Pool name** | **Composition** | **n** | **Rationale** |
| --- | --- | --- | --- |
| BalkanCore | Bulgaria_EIA + Greek IA/LBA + N. Macedonia_IA | 44 | Broad Balkan Iron Age substrate; maximizes temporal and geographic coverage |
| AnatoliaMax | Aegean + Central + Eastern + Southeastern Anatolian populations | 37 | Pan-Anatolian pool; captures full Anatolian ancestry axis |
| Mugla3 | Mugla_Camandras_Dalagoz + Mugla_Stratonikeia + Mugla_Degirmendere (Roman/Byzantine) | 23 | Aegean Anatolia specifically; closest geographic proxy to Anatolian ancestry in Gothic targets |
| Mugla4 | Mugla3 + Mugla_Samantas (Byzantine) | 29 | Sensitivity: adding later Byzantine component |
| BalkanCore + Mugla3 (union-south) | BalkanCore + Mugla3 combined | 67 | Union-south stress test; tests two-way model violation |
| BalkanCore + AnatoliaMax (union-south) | BalkanCore + AnatoliaMax combined | 81 | Union-south stress test; tests two-way model violation |
| Bulgaria_EIA (single pop.) | Bulgaria_EIA alone | 14 | Original baseline; small sample size comparison |

The convertf PACKEDANCESTRYMAP subset retained 359 individuals across 61 populations and 1,196,712 SNPs. Pooled labels were created through sequential relabelling of .ind files while keeping genotype and SNP files constant. Several early failures were traced to incorrect pooling order (resulting in “admixing pop ... has no samples” errors); these were corrected by building pooled .ind variants sequentially.

### S3.3 Complete Results: Sensitivity Analysis

#### *S3.3.1 KalideN (n=12): southern proxy sensitivity*

With the northern source fixed as Pol3 (Wielbark), three single-south proxies were tested. All produced highly significant signals (Z>4.5) with consistent timing estimates:

| **South proxy** | **Mean (gen)** | **SE** | **Z** | **nrmsd** |
| --- | --- | --- | --- | --- |
| BalkanCore (n=44) | 12.09 | 2.61 | 4.63 | 0.318 |
| Mugla3 (n=23) | 12.74 | 2.70 | 4.73 | 0.191 |
| AnatoliaMax (n=37) | 11.01 | 2.42 | 4.55 | 0.242 |

*Table S3.1. KalideN DATES results with Pol3 (Wielbark) as northern source. Timing estimates differ by less than two generations across all three southern proxies, well within the standard error of ~2.5 generations. Mugla3 produces the best fit quality (lowest nrmsd); AnatoliaMax yields the youngest point estimate.*

For comparison, when Bulgaria_EIA alone (n=14) was used as the southern source—the configuration tested in preliminary analyses—the result was 11.33±11.91 gen (Z=0.95): a similar point estimate but poor precision due to the small sample size. The BalkanCore pool, which adds Greek IA/LBA and North Macedonian IA populations to Bulgaria_EIA, improved the Z-score nearly fivefold (0.95→4.63) and reduced the standard error by 78% (11.91→2.61). This suggests that sample size and geographic breadth in the southern reference are critical for stable inference, while the underlying timing signal is robust.

#### *S3.3.2 KalideN: northern proxy sensitivity and signal specificity*

Adding Scandinavian Iron Age samples to the northern reference panel systematically shifted admixture dates older and inflated standard errors:

| **North proxy** | **South** | **Mean (gen)** | **SE** | **Z** | **nrmsd** |
| --- | --- | --- | --- | --- | --- |
| Pol3 (n=68) | BalkanCore | 12.09 | 2.61 | 4.63 | 0.318 |
| Pol3+Scandinavia (n=82) | BalkanCore | 14.63 | 4.59 | 3.19 | 0.311 |
| Pol3 (n=68) | Mugla3 | 12.74 | 2.70 | 4.73 | 0.191 |
| Pol3+Scandinavia (n=82) | Mugla3 | 17.26 | 4.63 | 3.73 | 0.170 |
| Pol3 (n=68) | AnatoliaMax | 11.01 | 2.42 | 4.55 | 0.242 |
| Pol3+Scandinavia (n=82) | AnatoliaMax | 13.60 | 3.37 | 4.04 | 0.230 |

*Table S3.2. Effect of adding Scandinavian IA to the northern reference. Adding Scandinavia shifts dates older by 2.5–4.5 generations and inflates SE by 40–76% across all southern proxies.*

Decomposition of the expanded-North effect reveals that Denmark_IA is the primary driver of the upward shift: Pol3+Denmark yields 16.42±3.53 gen (nrmsd=0.175), while Pol3+Sweden/Gotland yields 13.85±4.86 gen (nrmsd=0.195). Scandinavia alone (n=14, no Wielbark) produces 15.95±5.57 gen. The pattern is consistent with the detected admixture signal being specifically associated with Polish Wielbark-like ancestry rather than generic northern European Iron Age ancestry: Scandinavian proxies introduce a distinct allele-frequency contrast that possibly maps onto deeper population structure—potentially reflecting Proto-Germanic ethnogenesis (the Corded Ware → Scandinavia → Jutland → Wielbark migration trajectory), rather than the specific Gothic-associated admixture event detected at ~12 generations with Wielbark sources.

#### *S3.3.3 Pooled target KalideN+AKO (n=23)*

Pooling KalideN and AKO into a single target increased statistical power, yielding the strongest overall signal (Z=5.26 for AnatoliaMax). Timing estimates remained stable for BalkanCore and AnatoliaMax (~12 gen) and shifted slightly older for Mugla3 (~14.7 gen), possibly because AKO carries a somewhat different southern mixture relative to Mugla3 specifically:

| **South proxy** | **Mean (gen)** | **SE** | **Z** | **nrmsd** |
| --- | --- | --- | --- | --- |
| BalkanCore (n=44) | 12.12 | 2.76 | 4.39 | 0.221 |
| Mugla3 (n=23) | 14.65 | 3.14 | 4.67 | 0.160 |
| AnatoliaMax (n=37) | 12.44 | 2.36 | 5.26 | 0.186 |

*Table S3.3. KalideN+AKO pooled target with Pol3 northern source. The best overall model (AnatoliaMax, Z=5.26) is used as the primary result in the main text (Table 4).*

### S3.4 Pre-Mixed Substrate vs. Two-Pulse Model

The convergence of admixture timing across geographically and chronologically distinct southern proxies (§S3.3.1, S3.3.3) raises a question with direct historical implications: did the Balkan and Anatolian components of the southern substrate enter the Gothic gene pool as a single pre-mixed population, or as temporally separable admixture pulses? These models generate opposing predictions that can be formally tested against the DATES results.

#### *S3.4.1 Predictions*

**Pre-mixed substrate model.** If Balkan and Anatolian ancestries were already blended into a single demographic unit before encountering Wielbark-related groups, the following predictions hold: (1) Balkan-only and Anatolia-only southern sources should yield similar admixture dates, because both capture the same underlying event. (2) The two source types should be interchangeable as proxies, producing comparable Z-scores. (3) Pooling them into a union-south reference should degrade inference, because internal heterogeneity within the combined panel violates the two-way admixture assumption of DATES. (4) Admixture dates should be old, predating Gothic arrival in Bulgaria, because the mixing event occurred before the population encountered the substrate.

**Two-pulse model.** If Thracian/Balkan and Anatolian ancestries entered the Gothic gene pool as temporally separable events - for instance, Balkan admixture during Chernyakhov formation and Anatolian admixture after settlement in the eastern Roman Empire, the predictions reverse: (1) Balkan-only and Anatolia-only sources should yield different dates, capturing distinct events. (2) Each should have a unique admixture signature. (3) A union-south reference might capture both signals and thus improve or at least not degrade inference.

#### *S3.4.2 Results*

All four predictions of the pre-mixed model are confirmed by the data.

***Prediction 1: indistinguishable timing.*** For KalideN × Pol3, BalkanCore yields 12.09±2.61 gen and AnatoliaMax yields 11.01±2.42 gen—a difference of 1.08 generations, less than half the standard error. For the pooled KalideN+AKO target, BalkanCore yields 12.12±2.76 gen and AnatoliaMax yields 12.44±2.36 gen—a difference of 0.32 generations. The three-way comparison including Mugla3 (12.74 gen for KalideN, 14.65 gen for KalideN+AKO) shows slightly older estimates but remains within 1–2 standard errors of the other proxies. DATES cannot distinguish the admixture timing captured by Balkan versus Anatolian southern references.

***Prediction 2: interchangeable proxies.*** Both BalkanCore and AnatoliaMax produce highly significant signals when used as the sole southern source. For KalideN: Z=4.63 (BalkanCore) versus Z=4.55 (AnatoliaMax). For KalideN+AKO: Z=4.39 versus Z=5.26. The Z-scores are comparable in magnitude, indicating that both source types capture a real admixture signal of similar strength. Neither is markedly superior to the other, as expected if they represent different axes of the same blended substrate.

***Prediction 3: union-south degradation.*** Pooling Balkan and Anatolian sources into union-south references consistently degrades inference. For KalideN+AKO × Pol3:

| **South configuration** | **Mean (gen)** | **SE** | **Z** |
| --- | --- | --- | --- |
| BalkanCore alone | 12.12 | 2.76 | 4.39 |
| AnatoliaMax alone | 12.44 | 2.36 | 5.26 |
| BalkanCore + Mugla3 (union) | 15.54 | 5.12 | 3.04 |
| BalkanCore + AnatoliaMax (union) | 15.03 | 5.27 | 2.85 |

*Table S3.4. Union-south stress tests for KalideN+AKO × Pol3. Union-south configurations shift the point estimate 3 generations older, nearly double the standard error, and reduce Z-scores from ~4–5 to ~3. The pattern is reproduced for KalideN alone (not shown; comparable degradation).*

The degradation is consistent with the union-south reference introducing within-source heterogeneity that violates the two-way admixture assumption. When Balkan and Anatolian populations—which retain distinct allele frequency profiles despite being temporally co-introduced into the Gothic gene pool—are forced into a single reference panel, the resulting multi-modal frequency distribution reduces the effective contrast with the northern source and produces a noisier covariance decay.

***Prediction 4: old dates.*** All well-fitting models place admixture at ~11–13 generations before burial, corresponding to the 1st–2nd century CE for burial dates of ~360 CE (Aquae Calidae) and ~460 CE (AKO). This is centuries before Gothic settlement in Bulgaria (~376–382 CE) and predates even the earliest historically recorded Gothic intrusions into the eastern Roman Empire (~170 CE during the Marcomannic Wars). There is no signal of a recent post-arrival Balkan pulse, which would require dates of ~0–5 generations. The admixture event thus occurred mostly outside Bulgaria, at a time and place where Wielbark-related groups could encounter an already-blended Balkan-Anatolian population.

#### *S3.4.3 Summary*

Conversely, all three predictions of the two-pulse model fail: Balkan and Anatolian sources do not produce distinguishable dates, union-south pooling makes results worse rather than better, and there is no signal of a younger post-arrival pulse. These results constitute strong evidence that the Gothic-associated populations at Aquae Calidae and AKO inherited their Balkan and Anatolian ancestry, at least initially, from the same admixture event—mixing with a population in which these components were already blended—rather than from two separate encounters with Thracian and Anatolian groups at different times. The historical and geographic implications of this finding are discussed in the main text (§3.1).

### S3.5 Site-Specific Proxy Preferences

#### *S3.5.1 AKO (n=11)*

AKO shows a different optimal northern proxy from KalideN. With Mugla_Stratonikeia as the southern source, Chernyakhov as northern source yields Z=4.38 (17.23±3.94 gen), whereas Pol3 (Wielbark) yields only Z=2.64 with AnatoliaMax. This reversal is interpretable in terms of allele-frequency contrast: AKO’s ancestry is more evenly distributed between northern and southern components than KalideN’s, reducing the contrast with unmixed Wielbark and thus degrading DATES power. Chernyakhov, which is itself partially admixed (Wielbark + southern), provides a better-matched reference for AKO’s intermediate position. When KalideN was tested as a southern proxy for AKO, the result failed (Z<1), ruling out an ancestor-descendant relationship between the two groups.

#### *S3.5.2 KalideS (n=5): heterogeneity control*

KalideS produced weaker DATES signals across all configurations (Z~2.7–3.2, nrmsd~0.37–0.39). Notably, pooling the northern source into Pol3 actually worsened the signal compared to single-site Kowalewko, a counterintuitive result that is diagnostic: KalideS individuals have minimal northern ancestry (~10–20% per qpAdm), so even small heterogeneity introduced by pooling multiple Wielbark sites dilutes the already-weak contrast. The poor KalideS performance is itself evidence for the ethnogenesis interpretation: these individuals share identical Gothic material culture with KalideN but have a fundamentally different ancestry profile, consistent with Anatolian-ancestry individuals who adopted Gothic cultural identity without substantial Germanic admixture.

### S3.6 Control Tests on Non-Gothic Populations

To assess whether the ~12-generation admixture clock is specific to Gothic-associated targets, identical DATES contrasts were applied to Roman-period populations from outside Gothic archaeological contexts. Full results are reported in the main text (§2.3); here we note that Serbia_Viminacium_Roman_Rit (n=3; Olalde et al. 2023) failed all DATES contrasts (Z<1.8 with implausible estimates and nrmsd>0.66), and unpublished Roman-period Bulgarian populations (n=16 and n=3) similarly failed to produce stable admixture signals under Gothic-relevant contrasts. The ~12-generation Wielbark × Balkan/Anatolian signal is not a generic property of Late Antique Balkan populations but appears specifically in Gothic-associated targets.

### S3.7 Failed and Diagnostic Configurations

Several configurations produced impossible or implausible results. Rather than reflecting methodological error, these failures are informative: they identify proxy mismatches and violations of the single-pulse assumption that constrain the range of admixture models compatible with the data.

| **Configuration** | **Result** | **Diagnostic interpretation** |
| --- | --- | --- |
| AKO × Chernyakhov vs. AnatoliaReduced | Mean = 215.6 gen (implausible) | Chernyakhov and Anatolia reduced are not well-separated poles for AKO; the LD decay maps onto deep structure rather than a single pulse |
| AKO × BalkanCore vs. AnatoliaReduced | Mean = −51.1 gen (impossible) | BalkanCore and AnatoliaReduced do not bracket AKO ancestry; negative time indicates reversed covariance direction |
| KALIDE (full site) × Kowalewko vs. Samantas | Mean = negative (impossible) | Pooling KalideN+KalideS creates too much internal heterogeneity; the mixed target violates single-pulse assumption |
| KalideN+AKO × North_Full_Anchored vs. Mugla3 | Mean = 18.39±7.93 (Z=2.32, marginal) | Over-maxed North pool dilutes Wielbark-specific contrast; marginal significance despite large n indicates proxy mismatch |
| KalideS × Pol3 vs. various | Z = 2.7–3.2 (weak–marginal) | Insufficient northern component in KalideS (~10–20%) for robust DATES inference |

*Table S3.5. Selected failed and marginal configurations. Configurations producing implausible values (negative times, >200 generations) indicate that the specified reference populations do not bracket the target’s ancestry along the relevant axis, confirming that the ~12-generation signal detected with Pol3 × BalkanCore/AnatoliaMax/Mugla3 is specific to the Wielbark–Balkan/Anatolian contrast.*

In some runs, DATES produced runtime error messages but still generated complete output files; re-running produced identical results, indicating a software reporting issue rather than stochastic instability. Such runs are reported but interpreted cautiously.

### S3.8 Calendar Date Estimates

Calendar date estimates for all principal models are provided in Table 4 of the main text (§2.3), which serves as the authoritative reference for point estimates and 95% confidence intervals. Calendar conversion uses 29 years per generation; the burial date midpoint assumed for each target group is stated in Table 4. The sensitivity of calendar estimates to burial date assumptions is proportional to the uncertainty in the generation estimates themselves (SE ∼2.5 gen ≈ 73 years) and does not alter the primary conclusion that the admixture event predates Gothic settlement in Bulgaria.

### S3.9 Limitations

Several constraints affect the interpretation of these results. Target sample sizes remain modest (KalideN n=12, AKO n=11, KalideS n=5), limiting power for fine-grained within-site comparisons. Some source pools, particularly Chernyakhov (n=3), are small relative to the Balkan and Anatolian pools (n=37–44), potentially introducing stochastic effects. The Mugla proxies span Roman, Byzantine, and undated contexts; temporal offsets may affect interpretation, though the modest difference between Mugla3 and Mugla4 (adding Samantas Byzantine: 11.77 vs. 10.84 gen for KalideN) suggests limited sensitivity. DATES assumes a single admixture pulse; multi-pulse histories can produce composite signals that are difficult to decompose into individual events, and the broad confidence intervals (spanning ~250 years for the best model) preclude precise historical attribution. Generation time is assumed to be 29 years; alternative values would shift calendar interpretations proportionally. Weighted covariance decay curves for the principal models are provided as supplementary figures.
