## Supplemental Note S4 Archaeology Timeline for "Paleogenomic Evidence for Genetic Heterogeneity and Prior Admixture in Gothic-Associated Communities of Late Antique Bulgaria"

Supplementary Note S4: Historical Timeline and Archaeological Context

This note provides historical and archaeological context for the two Gothic-associated sites discussed in the study. Its purpose is to document chronology, site setting, and the basis for Gothic-associated attribution while distinguishing between archaeological observations, proposals made in the literature, and broader historical interpretations.

### Historical Timeline

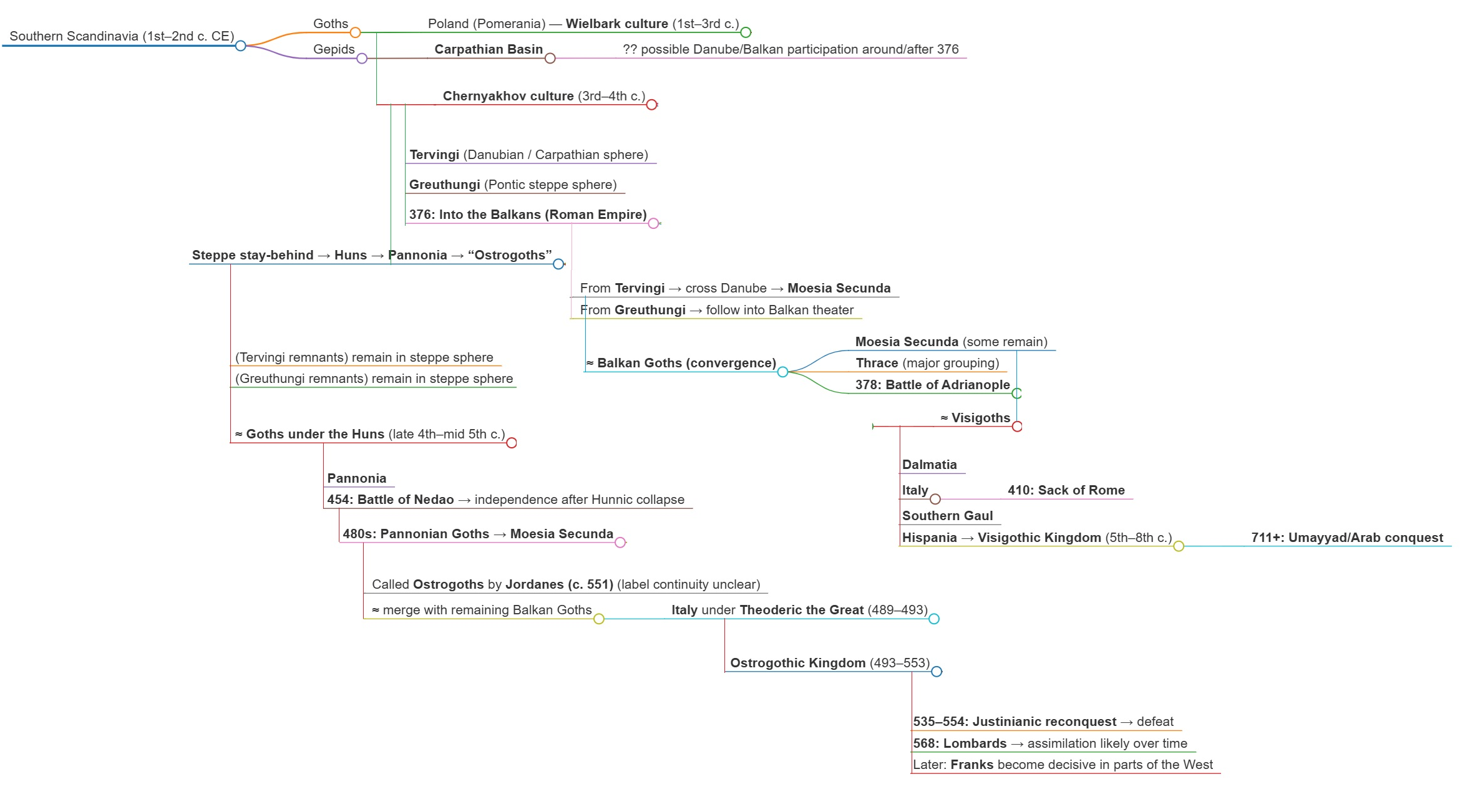

**Diagram S1.** Schematic flowchart of the historical development of Gothic groups from southern Scandinavia to their later Visigothic and Ostrogothic successor polities..

**Wielbark Culture (1st–4th centuries CE).** Emerging in northern Poland along the Vistula River and Baltic coast, the Wielbark culture represents the archaeological horizon most closely associated with early Gothic groups in historical reconstructions. It is characterized by material links to the southern Baltic zone and, in recent paleogenomic work, by affinities to Scandinavian-related ancestry profiles. In this study, Wielbark-associated populations serve as northern proxies in ancestry modeling and admixture dating, rather than as direct stand-ins for any single historically attested Gothic group.

**Chernyakhov Culture (late 2nd–5th centuries CE).** Formed in present-day Ukraine, Moldova, and Romania through interaction among populations of different cultural and biological backgrounds, the Chernyakhov horizon is generally understood as multi-ethnic. Its archaeological variability and the genetic substructure observed in recent ancient-DNA studies are consistent with a setting in which Wielbark-related groups interacted with local Dacian, Sarmatian, Alan, and other populations. In the present study, Chernyakhov-related populations are used as one class of northern/eastern proxy relevant to the formation of later Gothic-associated groups in the Balkans.

**3rd Century Raids (238–270s CE).** Gothic raids on Roman territory in modern Bulgarian lands began in 238 CE with an attack on Histria at the Danube mouth and escalated into sustained large-scale operations. The most consequential was Cniva’s coalition campaign of 249–251 CE, which crossed the Danube at Oescus (modern Gigen), defeated Roman forces at Nicopolis ad Istrum (250 CE) and at Beroe/Augusta Traiana (modern Stara Zagora, 250 CE), sacked Philippopolis (modern Plovdiv) after an extended siege with very large casualties, and culminated in the Battle of Abritus (251 CE, near modern Razgrad in Moesia Inferior) where Emperor Decius was killed — the first Roman emperor to fall in battle against a foreign enemy. The Gothic withdrawal was negotiated and involved retention of captives and spoils; the campaign was one of temporary military occupation, not permanent settlement. Maritime operations during 250–270 CE extended Gothic raiding to Pontus, Bithynia, Propontis, and the Aegean coast of Asia Minor, including a raid on Ephesus (263 CE) and documented capture of provincial populations in Cappadocia (c. 264/267 CE). These captives — among them, according to later tradition, the ancestors of Bishop Ulfilas — provide one historically plausible route by which Anatolian-related ancestry could have entered populations later associated with Gothic contexts, although the genetic data do not independently identify a single pathway or event. A combined Gothic-Herul invasion of 267–270 CE extended raiding through Macedonia and deep into Greece, and was eventually checked by Emperor Claudius II at the Battle of Naissus (268–269 CE, modern Niš, Serbia). The hot springs at Aquae Calidae near Anchialos (modern Burgas region) are specifically mentioned by Jordanes in connection with a Gothic force resting there after raiding activity in the later 3rd century, providing a direct historical association between the site and Gothic presence before any permanent settlement.

**4th Century Treaties and Settlements (332–382 CE).** The treaty of 332 CE between Constantine and the Tervingi Goths formalized relations along the lower Danube, granting trading rights and establishing Goths as foederati in the former Dacian buffer zone. Ulfilas was consecrated as Gothic bishop c. 341 CE and, following persecution by the Tervingian iudex Athanaric, led a community of Arian Christian Goths south of the Danube to settle near Nicopolis ad Istrum in mountainous Moesia Inferior (348 CE); this community, the “Minores Gothi,” persisted for at least two centuries and is the probable ancestral horizon for the Gothic ecclesiastical center at AKO. Valens’ Gothic war (367–369 CE) ended without settlement south of the river. The Hunnic advance of c. 370–375 CE collapsed the Chernyakhov culture and triggered the mass Danube crossings of 376 CE: the Tervingi under Fritigern crossed at Durostorum (modern Silistra, Bulgaria), followed by unauthorized crossings of Greuthungi, Alans, and Sarmatians. This combined force, ethnically already diverse, defeated and killed Emperor Valens at the Battle of Adrianople (9 August 378 CE, near modern Edirne on the Bulgarian-Turkish border). The subsequent Treaty of 3 October 382 CE under Theodosius I granted unprecedented autonomous settlement in Lower Moesia and Thrace, with the federated communities retaining arms, leadership, and internal governance in exchange for military service. This treaty established the primary, long-term Gothic presence in Bulgarian lands; the communities settled under it were the foundation population for subsequent phases documented at both sites in this study.

**5th Century Post-Hunnic Reconfigurations (453–488 CE).** After Attila’s death (453 CE) and the Battle of Nedao (454 CE), Pannonian Gothic groups under Amal leadership settled in Pannonia as Eastern Roman foederati (455–456 CE), where they had lived alongside Alans, Sarmatians, and former Hunnic subjects for two decades — the period during which some East Asian genetic components entered Gothic-associated populations. Simultaneously, the Thracian Goths — descendants of the 382 CE settlements — maintained a distinct, “deeply rooted” presence in Thrace and Moesia, receiving their own Roman subsidies. Famine conditions drove Pannonian Ostrogoths under Theodemir (father of Theodoric) to migrate to Macedonia in 473–474 CE, creating two competing Gothic federate communities in Bulgarian lands. Theodoric Amal’s own Balkan career began with a campaign to Singidunum in 469–470 CE; he subsequently conducted raiding campaigns through southern Bulgaria and the Rhodopes (476–477 CE), advanced to Marcianopolis (modern Devnya, Bulgaria) in 478 CE, and was formally granted command over Dacia Ripensis and Moesia Inferior with the title magister militum praesentalis in 483–484 CE. After eliminating his rival Theodoric Strabo’s heir in 484 CE, he united the Pannonian Ostrogoths and the long-established Thracian Gothic communities under a single leadership for the first time. Theodoric remained in Lower Moesia with his seat at Novae (modern Svistov) from c. 474 to 488 CE; in 487 CE his forces briefly threatened Constantinople itself. His departure for Italy in late 488 CE did not bring all Goths with him: significant populations remained in Moesia, Thrace, and Dacia Ripensis, forming the residual communities that continued in Bulgarian lands into the 6th century. The multiple churches and burial phases at AKO are consistent with occupation across this whole sequence of events, though archaeological evidence does not permit confident attribution of specific phases to named groups or events.

**6th–7th Centuries: Gradual Absorption.** Jordanes (writing c. 551 CE) describes Gothic communities in Moesia as “a numerous people, but poor and unwarlike,” subsisting on pastoralism and forestry near Nicopolis at the foot of the Haemus range — a description that matches the location of the AKO complex remarkably well and likely reflects the descendants of the long-established Moesian Gothic Christian community. Gothic soldiers and communities in various roles continued to serve the Eastern Roman state throughout the 6th century. Gradual absorption into Slavic and Byzantine populations occurred during the 6th–7th century Slavic migrations. The evidence from this later phase is better suited to documenting survival of cultural and ecclesiastical traditions than to reconstructing continuous biological descent, and the extent to which Gothic-associated genetic ancestry persists in modern Balkan populations remains beyond the scope of the present study.

### Archaeological and Genetic Context of Gothic Presence in the Balkans

The expansion of Hunnic power in the northern Black Sea region in the 4th century initiated a period of major political and demographic change across southeastern Europe. The Gothic groups visible in written and archaeological sources during the 4th–6th centuries were not historically static and should not be treated as a single uniform population. Accordingly, the present study does not use archaeology to prove a biologically coherent Gothic people; instead, it uses archaeology to define site context, chronology, and the grounds on which the sampled assemblages can be discussed as Gothic-associated.

Historical scholarship has long emphasized both continuity and diversity among Gothic groups in the Balkans. The archaeological record of the lower Danube and Thrace similarly points to repeated movement, settlement, interaction, and local incorporation. The two sites analyzed here—Aquae Calidae near Burgas and the complex at Khan Krum (AKO) near Shumen—are therefore best understood as contexts within a broader Gothic-associated historical horizon rather than as self-evident ethnic isolates.

#### Aquae Calidae (Akva Kalide)

The early Christian necropolis at Aquae Calidae near present-day Burgas was excavated in 2012. The cemetery occupies part of the abandoned Roman bath complex and presently comprises limited number of excavated graves, likely representing only a portion of a larger burial ground. Jordanes mentions the thermal springs of Anchialos in connection with Gothic raiders in the later 3rd century, providing a broad historical association for Gothic activity in the area. The excavated graves are broadly Christian in character, with predominantly east–west orientation, and are dated by the associated finds—coins, ceramics, and other grave goods—to the 4th century, most likely the third quarter of that century and not later than AD 375 (Momchilov and Klasnakov 2020).

The archaeological basis for describing the necropolis as Gothic-associated rests on a cluster of features rather than on any single diagnostic item. These include polished ceramic vessels with lattice-like ornament comparable to material from other Bulgarian sites with Gothic associations; cases of artificial cranial deformation similar to those documented at Kabile, Khan Krum, and Varna; the use of pottery tiles in some graves, a feature paralleled at sites such as Novae and Augusta Traiana; glass beads with parallels in other Bulgarian Gothic-associated burials; and a bone comb with a semicircular handle and concentric-circle ornament found close to the graves. Taken together, these observations support interpretation of the cemetery as part of a Gothic-associated mortuary horizon in the region.

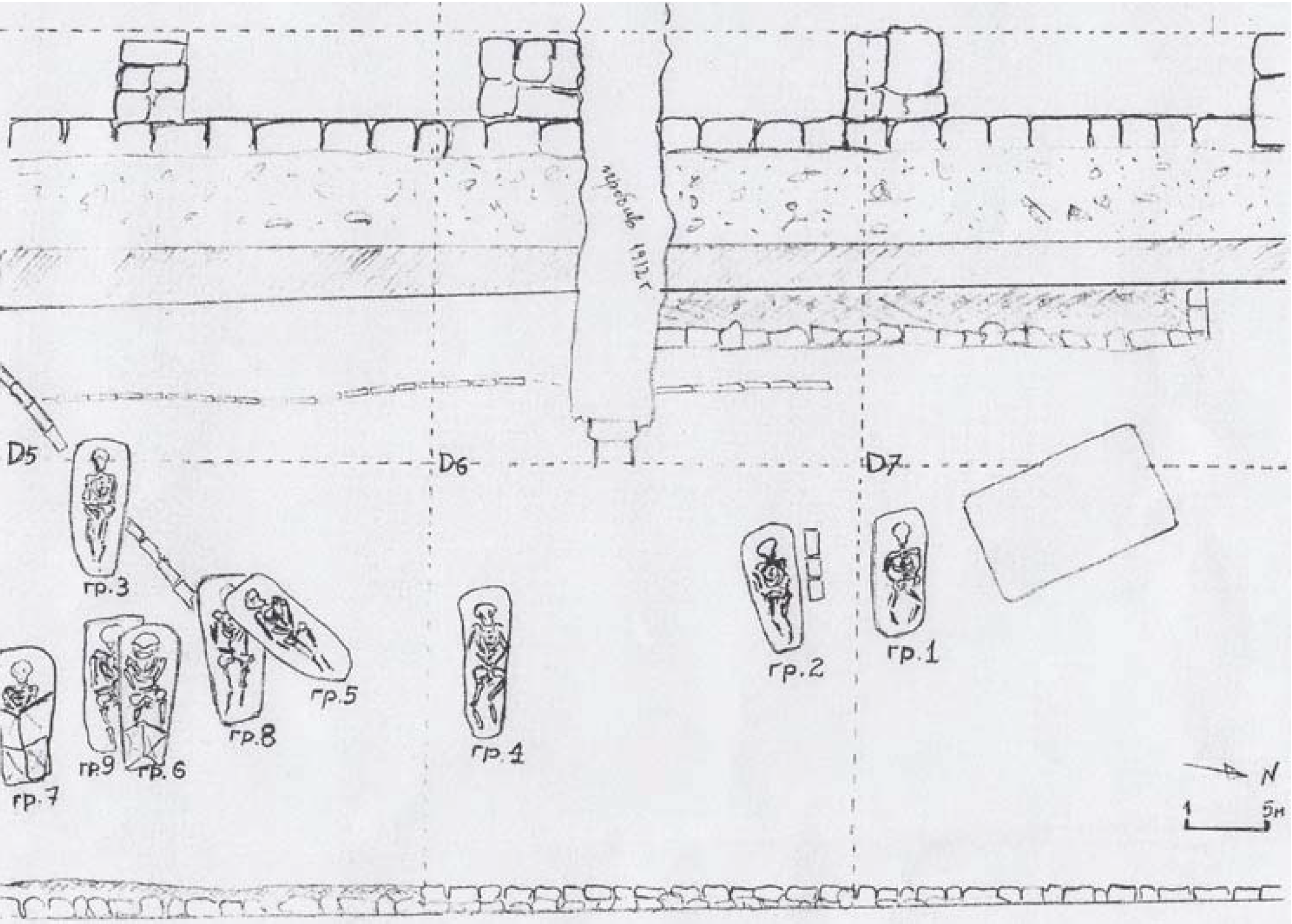

**Figure S1.** Archaeological plan of the early Christian necropolis at Aquae Calidae (Akva Kalide), near Burgas, Bulgaria. The burial area lies within the abandoned Roman bath complex, and most graves are oriented east–west. The plan documents reuse of the bath structures as funerary space. After Momchilov and Klasnakov 2020, p. 35.

Because the excavated cemetery is incomplete and the associated finds derive from a limited portion of the site, the social composition of the buried population cannot be reconstructed securely from archaeology alone. Considering the yet-to-be-studied parts of the necropolis and the archaeological material found in its vicinity, particularly the coins, the site was plausibly used as a burial ground for members of a local mixed community with Gothic associations, settled in the area of the major town of Anchialos sometime after the treaty of AD 332 between Constantine the Great and the Goths, and living here for at least two or three decades until the conflict of AD 376 forced the group to relocate. The individuals likely came from the northern frontier (Danube limes), and some may have married into the local population. It is possible also that some originated from mixed marriages between Gothic-associated groups and the local Byzantine population, or were descendants of provincial captives taken during the raids of the second half of the 3rd century. Such mixed origins are attested historically: Bishop Ulfilas, born in the Gothic lands north of the Danube around 311 CE, partially descended from Roman prisoners captured in a Gothic raid at Sadagolthina in Asia Minor (Wolfram 1988). The absence of children among the discovered burials may indicate that the Gothic members of this community came when already adults, not as a complete tribe, and subsequently settled and married within the Roman-era population present in the area — a pattern consistent with individuals serving the empire, possibly as a garrison to a nearby fortress. As an alternative scenario, the group may represent Arian Christian refugees who fled north of the Danube after the persecutions of AD 348, when Bishop Ulfilas and many followers relocated to Roman-controlled territory south of the river. The genomic results presented in the main text are compatible with all of these scenarios and support a population of heterogeneous biological origins participating in a shared Gothic-associated mortuary horizon, but the archaeological record by itself does not determine legal status, place of origin, or the precise mechanism of community formation.

#### Aul of Khan Omurtag (AKO), Khan Krum

The second site examined in this study is the late antique and early medieval complex near the modern village of Khan Krum, approximately 10 km from Shumen. Excavated since 1958, the site includes multiple occupation phases and five churches, four of them attributable to Late Antiquity and one to the 9th century. The modern name of the site refers to the fortified extra-capital royal residence of the Bulgarian rulers from the pagan period of the First Bulgarian Kingdom, built on the same place. It has long been discussed in Bulgarian scholarship as one of the most important Gothic-associated sites in present-day Bulgaria.

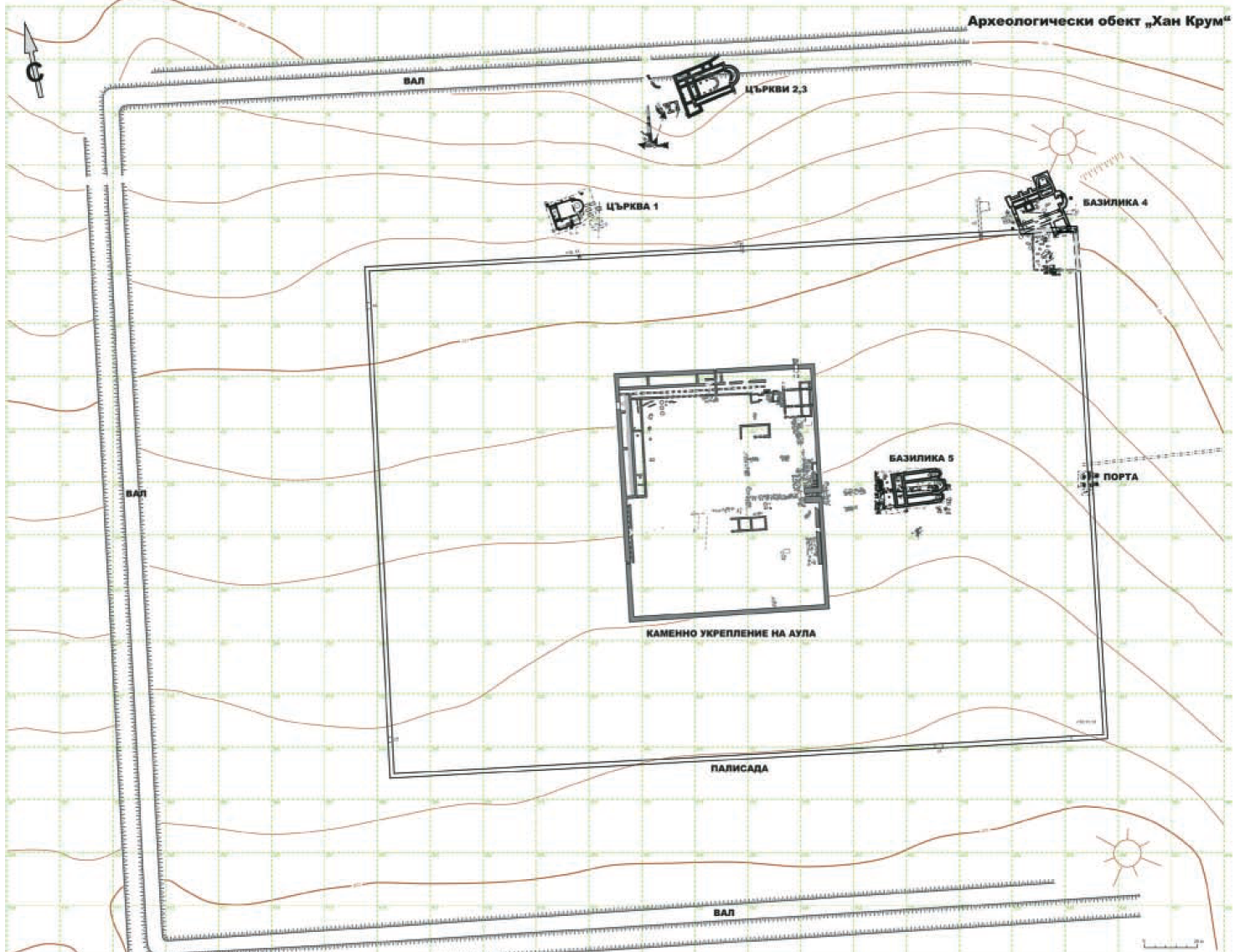

**Figure S2.** Site plan of the complex at the Aul of Khan Omurtag (AKO), near Shumen, Bulgaria, showing the late antique churches (N1–N4) and associated necropoleis (C1–C5). The complex preserves multiple Christian buildings and burial areas spanning Late Antiquity, with a later medieval Bulgarian church (N5) constructed over earlier remains. After Stoeva 2014.

**Church N1.** N1 is a relatively small one-nave basilica with an altar barrier characteristic of early Christian architecture. Coins recovered from the church and its surroundings range from the mid-4th to the mid-5th century, indicating use over a substantial period. The burning of the structure has been connected by excavators with regional disturbances in the early or mid-5th century, although an exact historical cause cannot be assigned with certainty. Associated grave goods, including a silver fibula, beads, and metal instruments from a grave in the narthex, are broadly consistent with a late 4th- to early 5th-century date.

**Church N2.** N2 is a large three-aisled basilica with an apse containing a synthronon, an architectural feature compatible with episcopal use. It has been dated to the later 5th or early 6th century. Burials excavated east of the church included rich grave goods of late antique elite character, indicating the presence of high-status individuals at the site.

**Church N3.** N3 is a one-aisled basilica underlying N2, showing that the complex developed through multiple building phases. Its plan has been compared with mid-4th-century church architecture elsewhere in Bulgaria, and coins from the relevant layer fall within the period AD 364–392. Fragments of high-quality wall painting were also recovered. Two sword depictions in the painted decoration have been interpreted by Balabanov as symbolically meaningful; however, the specific association of these motifs with Norse or Tervingian traditions remains a hypothesis rather than a settled identification. Church N3 does not have an independent necropolis; the burials located in its vicinity are attributed to the necropolis associated with Church N2 (Stoeva 2014).

**Church N4.** N4 was discovered beneath a large early medieval mound and is less securely dated because of disturbance by later activity and looting. On present evidence, its construction likely falls in the middle or second half of the 5th century. An octagonal structure with an apse was later erected on the same foundations. Disturbed rich burials within this area included individuals with artificial cranial deformation, traces of substantial bracelets, and fragments of chainmail, all consistent with a high-status funerary setting in the later history of the site.

**Church N5.** N5 is a 9th-century Bulgarian basilica resembling other churches built after the Christianization of the First Bulgarian State. Its construction over an earlier burial area indicates continued reuse of the site after the Late Antique phases relevant to the present study.

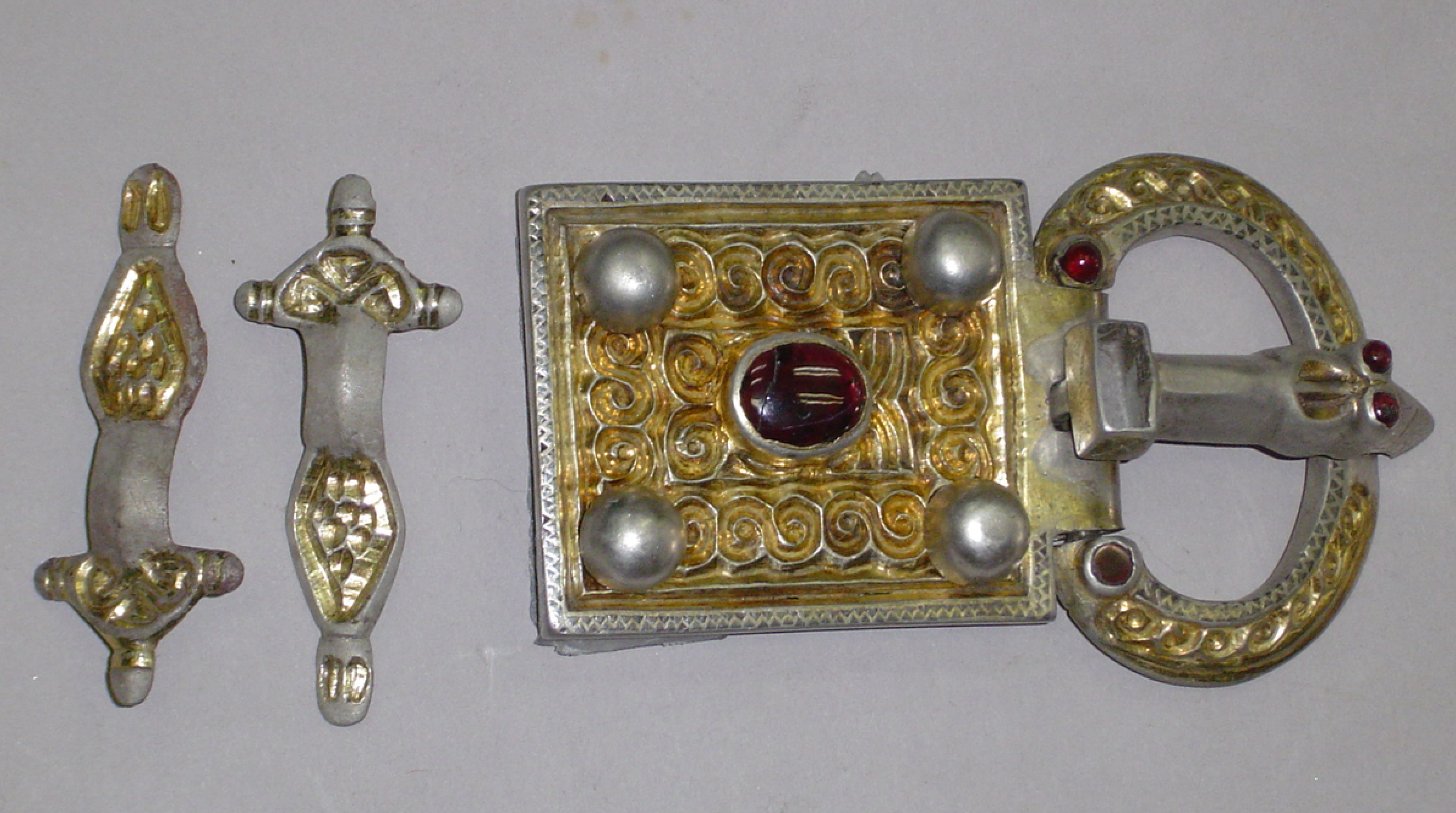

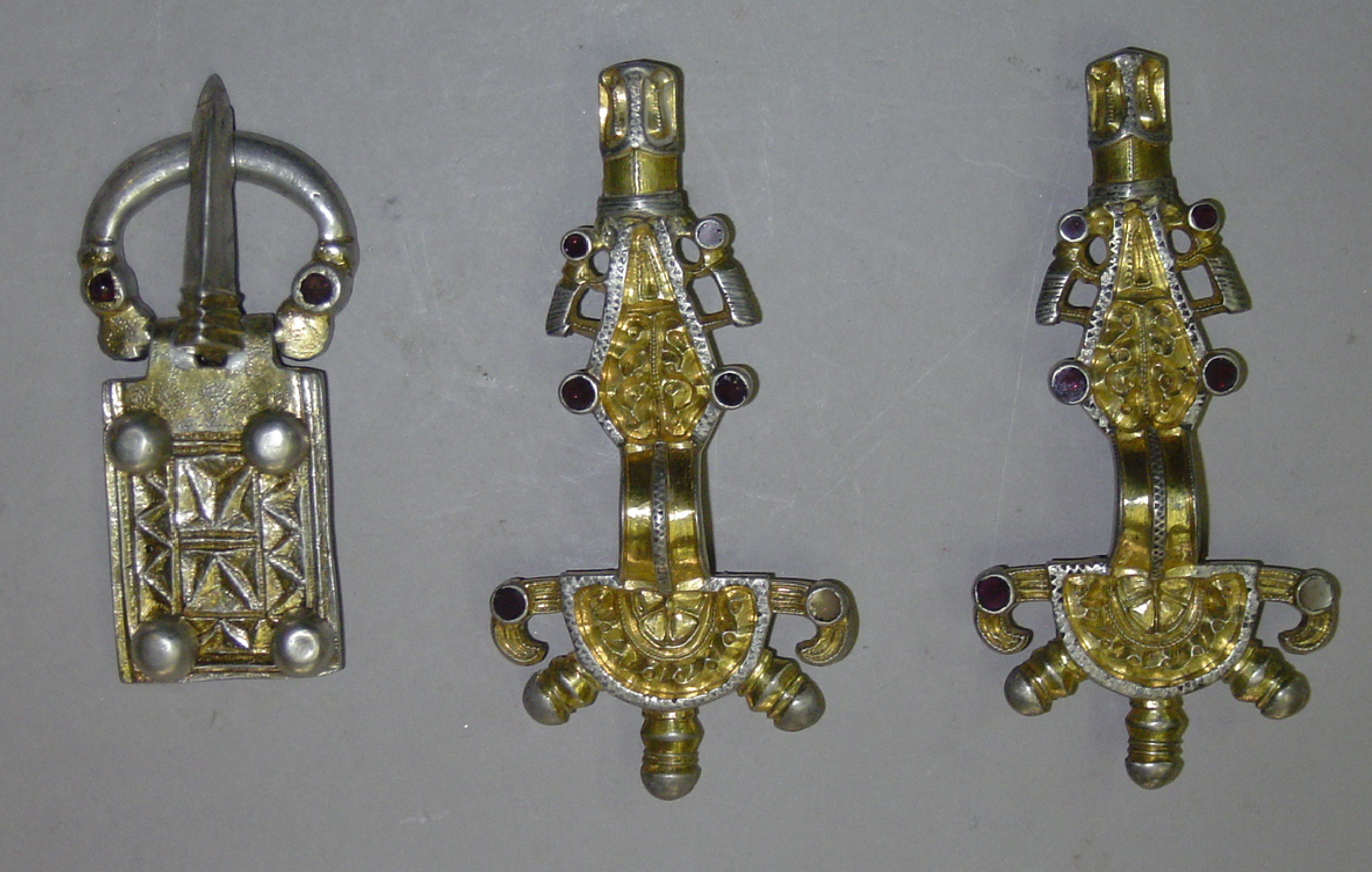

**Figure S3.** Grave goods from high-status burials at AKO Church N2. (A–B) Gold and gilded silver jewelry items from graves 1 and 2, displaying late antique elite styles characteristic of Migration Period assemblages. Such finds are consistent with the presence of high-status individuals within the Christian center and find parallels in other Gothic-associated elite burials. Images provided by the museum of Veliki Preslav.

The archaeological observations summarized above, together with written evidence for Gothic presence in northern Bulgaria, have led several Bulgarian researchers to interpret AKO as a Gothic bishopric center occupied from the mid-4th to the late 5th century and later reused in the Byzantine and Bulgarian periods. A possible connection with the wider ecclesiastical landscape associated with Ulfilas has often been proposed in the literature, although the precise localization of the communities described in the written sources remains uncertain. Jordanes’ reference to the 'Minor Goths' in Moesia therefore provides one possible contextual framework for discussion of the site, but not a direct identification.

Similarly, the multiple churches, changing burial phases, and high-status graves at AKO are consistent with long-term occupation by a substantial Christian community with Gothic associations. These features do not, however, determine whether individual phases should be linked to Ulfilas’ followers, later Ostrogothic groups, or other Gothic-affiliated populations in the region. In the present study, the archaeological evidence is used primarily to situate AKO within a Gothic-associated Christian horizon and to define its internal chronology, while the genomic analyses address questions of ancestry heterogeneity and demographic change that archaeology alone cannot resolve.
