## Supplemental Note S5 F3 Statistics for "Paleogenomic Evidence for Genetic Heterogeneity and Prior Admixture in Gothic-Associated Communities of Late Antique Bulgaria"

Supplementary Note S5: Outgroup f3 Affinity Statistics

*This note provides interpretation of the outgroup f3 statistics for the two Gothic-associated assemblages studied in the main text, complementing the full statistical output in Supplementary Table S1. The results are discussed in §2.1 of the main text; this note provides additional detail on the ranking structure and its relationship to the formal ancestry modeling.*

### S5.1 Analytical framework

The outgroup f3 statistic, here of the form f3(Mbuti; Target, Reference), measures shared genetic drift between a target population and a reference population relative to an outgroup (Mbuti). Higher f3 values indicate greater overall genetic affinity. The statistic is useful for characterizing broad affinity structure and for comparing relative similarity across sites and time periods; it does not by itself identify a direct source population or a migration route. Formal ancestry decomposition, source-model testing, and admixture dating are addressed in the main text through qpAdm, f4 statistics, PCA, and DATES (§2.1–2.3).

### S5.2 Overall affinity pattern

Both Gothic-associated assemblages show their highest f3 affinities to Central and Eastern European Migration Period populations rather than to eastern steppe or East Asian–shifted groups. The top-ranked comparators for both sites (f3 > 0.052, |Z| > 70) include Wielbark-associated and other Iron Age or Migration Period populations from Poland and Central Europe, as well as Danubian Roman and Late Antique Balkan comparators. This broad pattern is consistent with the ancestry inferences from PCA and qpAdm reported in the main text (§2.1–2.2).

### S5.3 Site-level contrast

The two sites show a consistent directional difference in their f3 profiles. AKO shows slightly higher affinity to several northern and central European comparators, including Wielbark-associated and other Migration Period “Germanic” groups, whereas Aquae Calidae shows relatively stronger affinity to Bulgaria_LAntiquity and other Balkan comparators. AKO's slightly stronger northern affinity and Aquae Calidae's stronger pull to Bulgaria_LAntiquity (f3=0.0524, rank #1 vs. rank #21 at AKO) are consistent with the qpAdm ancestry proportions reported in §2.2. At the same time, the numerical differences in f3 are modest, so they should be interpreted as shifts in relative affinity rather than as evidence that the two sites belonged to wholly separate population traditions.

### S5.4 Selected comparator classes

Several comparator groups warrant specific comment. Wielbark-associated populations and related Iron Age Polish groups rank among the strongest affinities for both sites, which is consistent with the manuscript’s use of Wielbark-like populations as one northern reference frame. Danubian and Central European Late Antique groups, including Hungary_Langobard and Austria_Klosterneuburg_Roman, also rank very highly; this reflects shared Central/Eastern European ancestry structure across the Migration Period world rather than direct historical contact with any specific group. Bulgaria_LAntiquity ranks especially highly for Aquae Calidae, which is compatible with the stronger Anatolian/southern ancestry component documented by qpAdm for that site (§2.2). By contrast, populations with strong East Asian–related ancestry, such as East Asian–shifted Hun groups, rank much lower for both sites. At the group level this supports the view that neither cemetery was dominated by an eastern steppe ancestry profile, although it does not exclude limited eastern input in particular individuals.

### S5.5 Relationship to formal ancestry modeling

The outgroup f3 results are complementary to the formal ancestry modeling in §2.2–2.3. Three points deserve note. First, both sites participate in a northern/central European affinity zone that includes Wielbark-related populations. Second, Aquae Calidae shows a somewhat stronger pull toward Late Antique and Balkan comparators than AKO, in line with its more southern-shifted qpAdm profile. Third, the fact that Chernyakhiv-related populations do not always rank among the very top f3 hits does not contradict their use in the manuscript: outgroup f3 summarizes overall similarity, whereas qpAdm and DATES test whether a population can serve as a statistically useful proxy within a specific admixture model. Because Chernyakhiv was itself heterogeneous, moderate rather than top-ranked f3 values are not unexpected.

### S5.6 Summary of affinity patterns by comparator class

Table S5-1 summarizes the broad affinity tendencies by comparator class. Full f3 statistics for all comparator populations are provided in Supplementary Table S1.

| **Comparator class** | **Illustrative examples** | **Pattern in current f3 results** | **Interpretation** |
| --- | --- | --- | --- |
| Northern/Central European high-affinity set | Wielbark; Germany_EarlyMedieval; Hassleben_Germanic | High or near-high f3 values for both sites; slightly stronger tendency at AKO | Supports a northern/central European affinity component, but does not by itself prove a single migration route or direct descent from any one named group. |
| Danubian / Migration Period comparators | Hungary_Langobard; Austria_Klosterneuburg_Roman; Slovakia_TesarkeMLynany_Germanic | Among the highest-ranked comparators for both sites | Compatible with shared ancestry structure across the wider Danubian and Migration Period world; should not be read as evidence of one-to-one historical continuity. |
| Late Antiquity / Balkan comparators | Bulgaria_LAntiquity; Slovenia_EIA; Bulgaria_EIA | Relatively stronger pull at Aquae Calidae than at AKO | Consistent with greater southern/local ancestry in Aquae Calidae, in agreement with the manuscript’s qpAdm results. |
| Chernyakhiv-related populations | Chernyakhiv_Culture and related northern/eastern proxies | Present at moderate affinity rather than always at the very top of the ranking | Not inconsistent with manuscript use: heterogeneity within Chernyakhiv and differences between descriptive similarity and formal admixture modeling can produce this pattern. |
| East Asian–shifted steppe groups | Hun_oEastAsian; Kazakhstan_Nomad_Hun_Sarmatian | Lowest or among the lowest f3 values for both sites | Supports the absence of a dominant East Asian–shifted ancestry profile at the cemetery level; does not exclude minor eastern input in isolated individuals. |

### S5.7 Limitations

Several interpretive limitations apply to these results. Small rank differences — many f3 values differ only at the third or fourth decimal place — should not be treated as strong historical signals. High affinity to a comparator does not establish direct ancestry from that population, particularly when targets are admixed and when comparison groups differ in date, sample size, and internal heterogeneity. Low ranking of a comparator such as Bulgaria_RomanPeriod.TW does not constitute a strong negative claim about population continuity; such patterns may reflect sample composition, chronological offset, and the fact that f3 captures whole-genome similarity rather than any one ancestry component. Outgroup f3 cannot determine whether cultural affiliation was reproduced through ecclesiastical, linguistic, legal, or other social mechanisms; those questions require integration with the archaeological and historical record.

### S5.8 Summary

The outgroup f3 results support a broad northern/central European affinity profile for both Gothic-associated assemblages, a stronger Balkan/Late Antique pull in Aquae Calidae compared with AKO, and uniformly low affinity to East Asian–shifted steppe groups. These patterns are descriptively consistent with the heterogeneity and differential site composition documented through PCA, qpAdm, f4 statistics, uniparental markers, and DATES, and they reinforce the conclusion that the two sites shared a Gothic-associated cultural horizon without being genetically identical. Full statistical results are in Supplementary Table S1.
